# Interspecies variation of larval locomotion kinematics in the genus *Drosophila* and its relation to habitat temperature

**DOI:** 10.1101/2020.11.06.371120

**Authors:** Yuji Matsuo, Akinao Nose, Hiroshi Kohsaka

## Abstract

Speed and trajectory of locomotion are characteristic traits of individual species. During evolution, locomotion kinematics is likely to have been tuned for survival in the habitats of each species. Although kinematics of locomotion is thought to be influenced by habitats, the quantitative relation between the kinematics and environmental factors has not been fully revealed. Here, we performed comparative analyses of larval locomotion in 11 *Drosophila* species. We found that larval locomotion kinematics are divergent among the species. The diversity is not correlated to the body length but is correlated instead to the habitat temperature of the species. Phylogenetic analyses using Bayesian inference suggest that the evolutionary rate of the kinematics is diverse among phylogenetic trees. The results of this study imply that the kinematics of larval locomotion has diverged in the evolutionary history of the genus *Drosophila* and evolved under the effects of the ambient temperature of habitats.

## Introduction

Kinematics of animal locomotion is a critical trait enabling each species to survive in their habitats (Alexander, 2006). Movement patterns have been sculpted during their evolution by adaptation to their environments, and could have diverged among species (Kappeler, 2010). Comparative analyses have identified several examples of differences in the kinematics of locomotion within a group of related species, including insects (Theunissen et al., 2015), reptiles (Fuller et al., 2011), birds (Stoessel & Fischer, 2012), and primates (Thompson et al., 2018). To take an example, two gecko species inhabiting either sandy or rocky environments have been shown to exhibit distinct postures (Fuller et al., 2011). Whereas the interspecies divergence in locomotion patterns can be observed in various phylogenetic branches of the animal kingdom, quantitative comparative analyses of locomotion kinematics remain limited.

Flies of the genus *Drosophila* have long been used as a model to study interspecific diversification and evolution (Markow & O’Grady, 2005). One salient example of the interspecific variations in the genus *Drosophila* is food for larvae. Some species eat multiple kinds of foods (“generalists” including *Drosophila melanogaster*) while others have strong preferences in food (“specialists” including *Drosophila sechellia*, which has specialized to feed on Morinda fruits) (Markow, 2015; Watada et al., 2020; Watanabe et al., 2019). These diverged species in the genus *Drosophila* offer an opportunity to perform interspecific comparisons in various animal traits including larval locomotion.

Fly larvae, or maggots, have been widely used in the study of the kinematics of locomotion (Berrigan & Lighton, 1993; Berrigan & Pepin, 1995; Fox et al., 2006; Green et al., 1983; Ruiz-Dubreuil et al., 1996; Sokolowski, 1985; Sokolowski et al., 1997). Among the fly species, *Drosophila melanogaster* (*Dmel*) is one of the most examined species, especially by virtue of the availability of resources in genetics and connectomics (Ohyama et al., 2015; Schneider-Mizell et al., 2016; Simpson & Looger, 2018; Venken et al., 2011). *Dmel* larvae locomote by a sequence of forward crawling, and changes of crawling direction achieved by bending their bodies (Green et al., 1983). The kinematics of larval locomotion is affected by ambient temperature (Kwon et al., 2010; Liu et al., 2003; Ohyama et al., 2013; Rosenzweig et al., 2005a). In thermotaxis behaviour in temperature gradient environments, larvae regulate the length of crawling runs between turns and the size and direction of turns (Luo et al., 2010), and the probability of turns is also affected by ambient temperature gradients (Klein et al., 2015). In contrast to the intensive studies on larval behavior in *Dmel*, locomotion kinematics in the larvae of its sister species in the genus *Drosophila* remains unclear.

Here we conducted an interspecies comparison of the kinematics of larval locomotion in the genus *Drosophila*. We address two questions in this study: are locomotion kinematics of larvae similar among *Drosophila* species?; and, if the kinematics are diverged, what factors are related to the diversity? To this aim, we recorded locomotion of larvae of 11 *Drosophila* species and extracted kinematic parameters using the tracking software FIMTrack (Risse et al., 2017). Clustering analysis with Jensen-Shannon divergence and statistical analyses show that two kinematics parameters (bend probability and crawling speed) differ among the *Drosophila* species. We found that kinematics varies with habitat temperature but not with body size. The relationship between the kinematics and minimum habitat temperature held at two distinct ambient temperatures: 24°C and 32°C. Phylogenetic analyses of these kinematics, based on Bayesian inference (Hohna et al., 2016), suggests that the rate of evolution of the kinematics is diverged among phylogenetic branches. Among eight traits we tested, the evolution of the crawling speed at 24°C and 32°C were correlated. Consequently, our results suggest that the kinematics of larval locomotion in the genus *Drosophila* diverged in response to environmental variation in ambient temperature.

## Results

### Kinematic analysis of crawling and bending behaviour in *Drosophila* larvae

In this study, we analyzed the kinematics of larval locomotion. To this aim, we recorded the locomotion of fly larvae that were crawling freely on an agarose substrate stage on a temperature-controlled plate (Figure 1A; see Method section for details). We placed eight to ten larvae at the centre of the stage, which was kept at 24°C, illuminated them with infrared light, which is invisible to fly larvae and does not affect their behaviour, and recorded larval locomotion for three minutes at five frames per second (Figure 1B). Maximum projection of the time-series images shows traces of larval locomotion as multiple curves of multiple larvae (Figure 1C). In an example of *Drosophila melanogaster* (Figure 1D), the traces show smooth curved lines interconnected with angles where larvae exhibit turning behaviour and change crawling direction, which is consistent with previous studies (Berni et al., 2012; Gershow et al., 2012; Gomez-Marin et al., 2011; Gomez-Marin & Louis, 2014; Green et al., 1983; Sims et al., 2019; Tastekin et al., 2015). Larvae under these conditions predominantly exhibit forward locomotion in which muscular contraction is propagating from the posterior to anterior segments. Characteristics of larval locomotion can be described by two measures: the bend angle and centroid speed (Gershow et al., 2012; Gomez-Marin et al., 2011). The bend angle measures the angle of body axis bending (Figure 1E) and the centroid speed is the speed of the position of the larval centroid (Figure 1F). To obtain these values for each larva at each time frame, we used the object tracking software for small animals, FIMTrack (Risse et al., 2017). The turning behaviour can be detected by the change in the angle of the body axis (Figure 1E) and the reduction in the centroid speed (Figure 1F), as reported previously (Gershow et al., 2012; Gomez-Marin et al., 2011). Accordingly, we used the bend angle and centroid speed for quantitative analysis of larval locomotion in this study.

**Figure 1.**
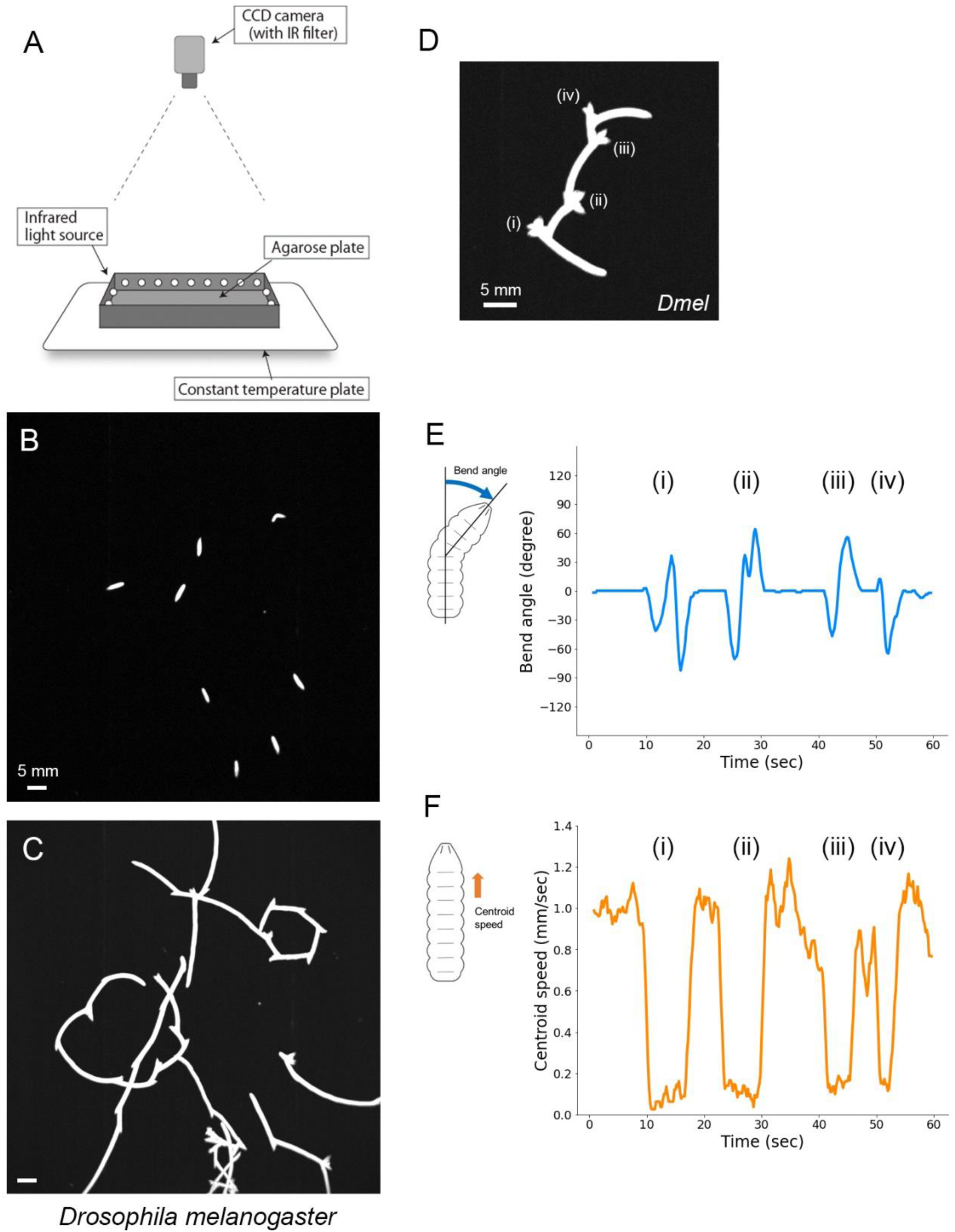
Measurement of larval crawling. (A) Setup of the recording of larval crawling with infrared light and temperature control plate. (B) An example image of multiple larvae recorded by the setup (A). (C) The trajectories of larval locomotion of *Drosophila melanogaster* recorded for two minutes. (D) A trajectory of single *Drosophila melanogaster* locomotion. Locations where the larva changes its direction are labelled (i) to (iv). (E and F). Bend angle (E) and centroid speed (F) of the larva in (D). Labels (i) to (iv) correspond to those in (D).

### Classification of locomotion properties in the genus *Drosophila*

For interspecies comparison of *Drosophila* larval locomotion, we collected 11 fly species in the genus *Drosophila*, whose genome sequences have been read (A. G. Clark et al., 2007; Miller et al., 2018), and for which living individuals are available from fly stock centres (KYORIN-Fly, Fly Stocks of Kyorin University; KYOTO Stock Center (DGRC) at the Kyoto Institute of Technology). The 11 *Drosophila* (*D.*, hereafter) species consist of *D. ananassae* (*Dana*)*, D. erecta* (*Dere*)*, D. mauritiana* (*Dmau*)*, D. melanogaster* (*Dmel*)*, D. mojavensis* (*Dmoj*)*, D. persimilis* (*Dper*)*, D. pseudoobscula* (*Dpse*)*, D. sechellia* (*Dsec*)*, D. virilis* (*Dvir*), *D. willistoni* (*Dwil*) *and D. yakuba* (*Dyak*). Nine species (all but *Dvir* and *Dmoj*) are classified as subgenus *Sophophora* and the remaining two (*Dvir* and *Dmoj*) are *non-Sophophora* species. We plotted the centroid speed and bend angle of freely crawling larvae of each species (Figure 2A). In all the cases, data points accumulated around a region where the bend angle is zero (Figure 2A), which reflects an observation that larvae do not bend during the majority of the time (Figure 1D). We noticed that the deviations from zero in bend angle axis are different among species. For example, data points in *Dvir* are scattered along the bend angle axis more than those in *Dwil* (Figure 2A). To quantify similarities in the distribution of the two-dimensional plots among the species, we calculated Jensen-Shannon divergence, which measures the similarity of two probability distributions (Lin, 1991). We classified the 11 species based on the similarity in the probability distribution of the crawling speed and bend angle plots by hierarchical clustering (Figure 2B and 2C). The classification analysis shows that four species (*Dmel, Dvir, Dmoj and Dper*) form a cluster. These four species show scattered data points along the bend angle axis compared with the other seven species (Figure 2A). This observation suggests that the kinematics of larvae in the genus *Drosophila* is diverse among species.

**Figure 2.**
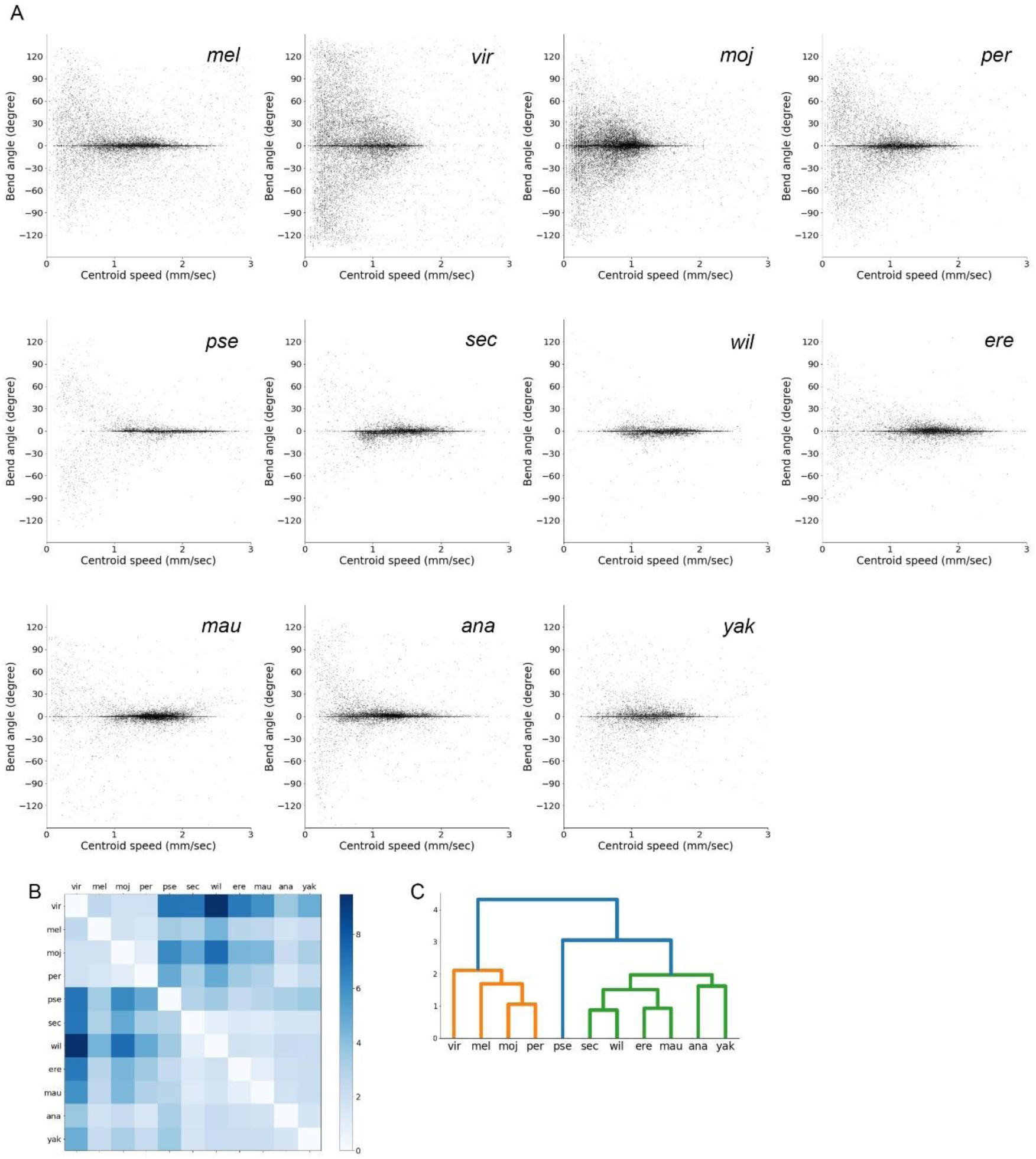
Interspecific comparison of kinematics of larval crawling in the genus *Drosophila*. (A) Plots of the speed of the centroid of larvae and the bend angle in the 11 *Drosophila* species. Each point corresponds to a datum of a single larva of a species in a single time frame. The number of larvae in each species is the following: *Dvir*: n=24; *Dmel*: n=24; *Dmoj*: n=30; *Dper*: n=27; *Dpse*: n=19; *Dsec*: n=21; *Dwil*: n=18; *Dere*: n=22; *Dmau*: n=26; *Dana*: n=22; *Dyak*: n=16. (B) Jensen-Shannon divergences of the probability density of (A). (C) Hierarchical clustering of the kinematics of larval locomotion of the 11 *Drosophila* species.

### Definition of the bend probability and crawling speed

The clustering analysis suggests that the kinematics is divergent among species. To interpret the diversity in terms of locomotion behaviour, we defined two indices: the bend probability and crawling speed. The bend probability measures how often larvae bend their body laterally. We set the minimum angle of bend as 20 degrees, which was used previously (Risse et al., 2013), and labelled larvae that bent more than this threshold angle to the right or left side as “Bending” (Figure 3A and 3B). This threshold allowed us to extract the difference in the bending rate among the species we examined, and we found that while *Dmel* larvae exhibit larger bend angles than this threshold (Figure 3A), the bend angles in *Dwil* larvae are mostly less than the threshold (Figure 3B). We defined the bend probability of each single larva as a ratio of the number of time frames labelled “Bending” to the total frame number. To define the second index crawling speed, we labelled larvae that bent less than the threshold as “Crawling” (Figure 3A and 3B). We defined the median of centroid speeds of larvae that were labelled “Crawling” as the crawling speed for each single larva. By these definitions, we calculated the bend probability and crawling speed of each larva of the species.

**Figure 3.**
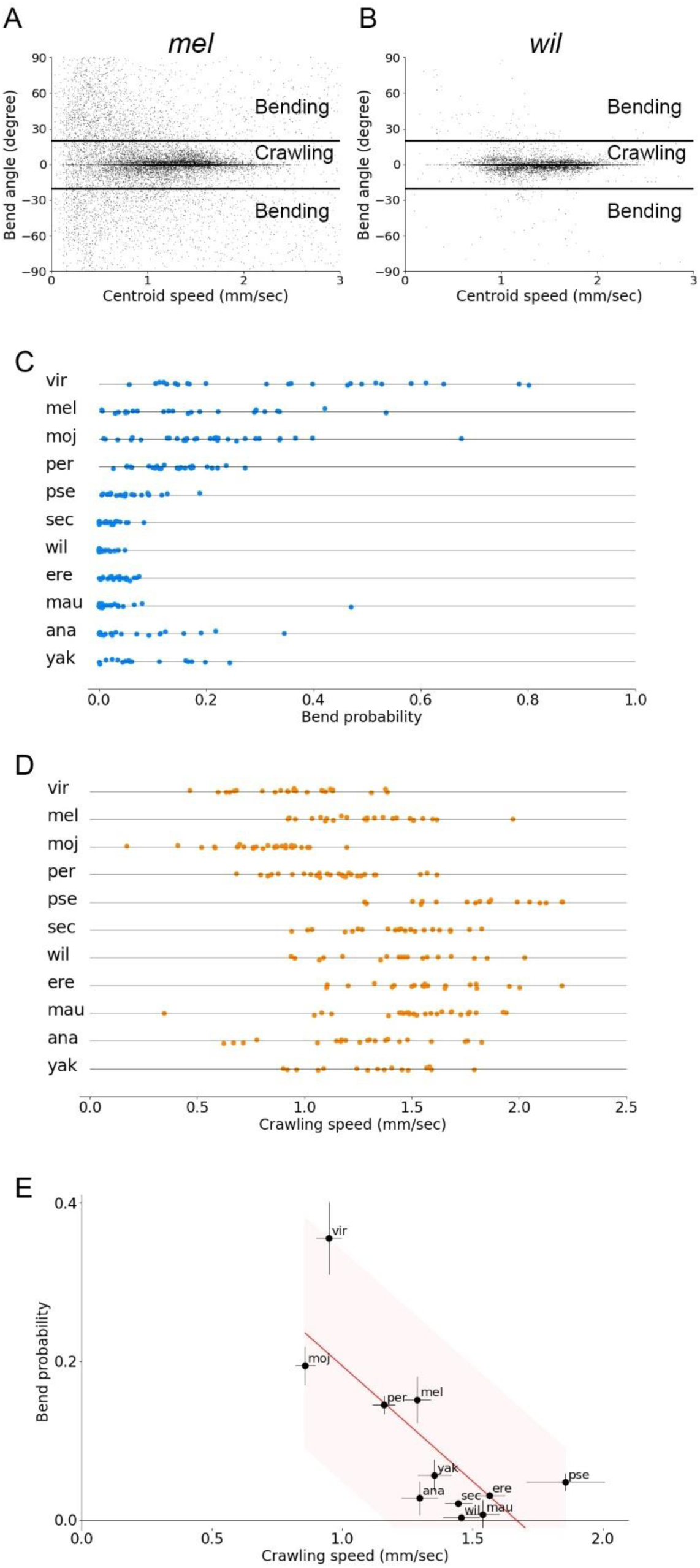
Bend probability and crawling speed of larval locomotion in the 11 *Drosophila* species at 24°C and their relationship to habitat temperature of each species. (A and B) Plots of the speed of the centroid of larvae and the bend angle of *D. melanogaster* (A) and *D. willistoni* (B). Each point corresponds to a datum of a single larva of a species at a single time frame. Thick horizontal lines denote the threshold between crawling and bending. (C) Bend probability of individual larvae of each species. (D) Crawling speed of individual larvae of each species. Sample numbers in (C) and (D) are the following: *Dvir*: n=24; *Dmel*: n=24; *Dmoj*: n=30; *Dper*: n=27; *Dpse*: n=19; *Dsec*: n=21; *Dwil*: n=18; *Dere*: n=22; *Dmau*: n=26; *Dana*: n=22; *Dyak*: n=16. (E) Scatter plot of bend probability at 24°C against crawling speed at 24°C. Median ± sem is shown. The red line shows the linear regression function and the shaded area represents the 95% confidence band. The point estimate of the Pearson correlation and its 95% confidence interval is -0.76 and [-0.94, -0.30].

### Kinematics of larval locomotion is diverse among the *Drosophila* species

We plotted the bend probability of each species (Figure 3C). A statistical analysis shows that the bend probability of the species is diverse (p = 6.8 x 10^-25^, Kruskal-Walls test). While *Dvir* larvae exhibited frequent bending (bend probability: 0.36 ± 0.05, n = 24), *Dwil* larvae rarely bend (bend probability: 0.003 ± 0.004, n = 18) (Figure 3C). We also plotted the crawling speed of the *Drosophila* species (Figure 3D). The statistical analysis shows that the speed is also diverse among the species (p = 9.3 x 10^-24^, Kruskal-Walls test). For example, *Dwil* larvae crawl faster than *Dvir* larvae do (crawling speed: 1.46 ± 0.07 mm/sec, n = 24 in *Dwil*; crawling speed: 0.95 ± 0.05 mm/sec, n = 18 in *Dvir*). These analyses indicate that the bend probability and crawling speed are differentiated in the genus *Drosophila*.

To capture the trends in the diversity of larval kinematics, we plotted the data in the space of the crawling speed and bend probability (Figure 3E). The graph shows a negative correlation between them (Pearson correlation = -0.76, p = 0.0064). While we used 20 degrees as the threshold for defining the bend probability (Figure 3A and 3B), the negative correlation between the crawling speed and bend probability is robust to the choice of the threshold (Pearson correlation = -0.80, p = 0.0030, when the threshold is 10 degrees; Pearson correlation = -0.71, p = 0.014, when the threshold is 30 degrees; Supplementary Figure 1A - 1C). Furthermore, the median of absolute values of bend angle instead of the bend probability exhibits a negative correlation to the crawling speed (Pearson correlation = -0.66, p = 0.026; Supplementary Figure 1D). To sum, the crawling speed and bend probability are diverse among the *Drosophila* species and negatively correlated.

### Comparison between intraspecies and interspecies variability

The comparison of the 11 *Drosophila* species represents the natural variability in larval locomotion across them. Here, it should be noted that we used a single strain for each species in the analysis. For that, the variability appeared above could result not only from interspecies but also intraspecies diversity. Even if there was no interspecies variability, sampling data from population with high intraspecies variation could lead to an apparent diversity across the species. To address this issue, we compared intraspecific deviation with interspecific variability. If intraspecific diversity is a dominant factor for the variability, the deviation within species should be comparable to that among species. In contrast, if interspecific diversity is a major cause, the deviation within species would be smaller than that across species. We examined intraspecies variation in two species in subgenus *Sophophora* (*Dmel* and *Dana*) and one *non-Sophophora* species (*Dvir*) (Figure 4). Two isofemale strains of each species were obtained from the Kyorin stock centre and the larval kinematics of them were measured. To evaluate the contribution of the interspecific deviations to the total deviations, we conducted the analysis of variance (ANOVA). We found a significant interspecific difference in the bend probability (p=0.008, the one-way ANOVA) although the difference in the crawling speed was marginal (p = 0.08, the one-way ANOVA). This observation implies the existence of interspecies diversity in larval locomotion.

**Figure 4.**
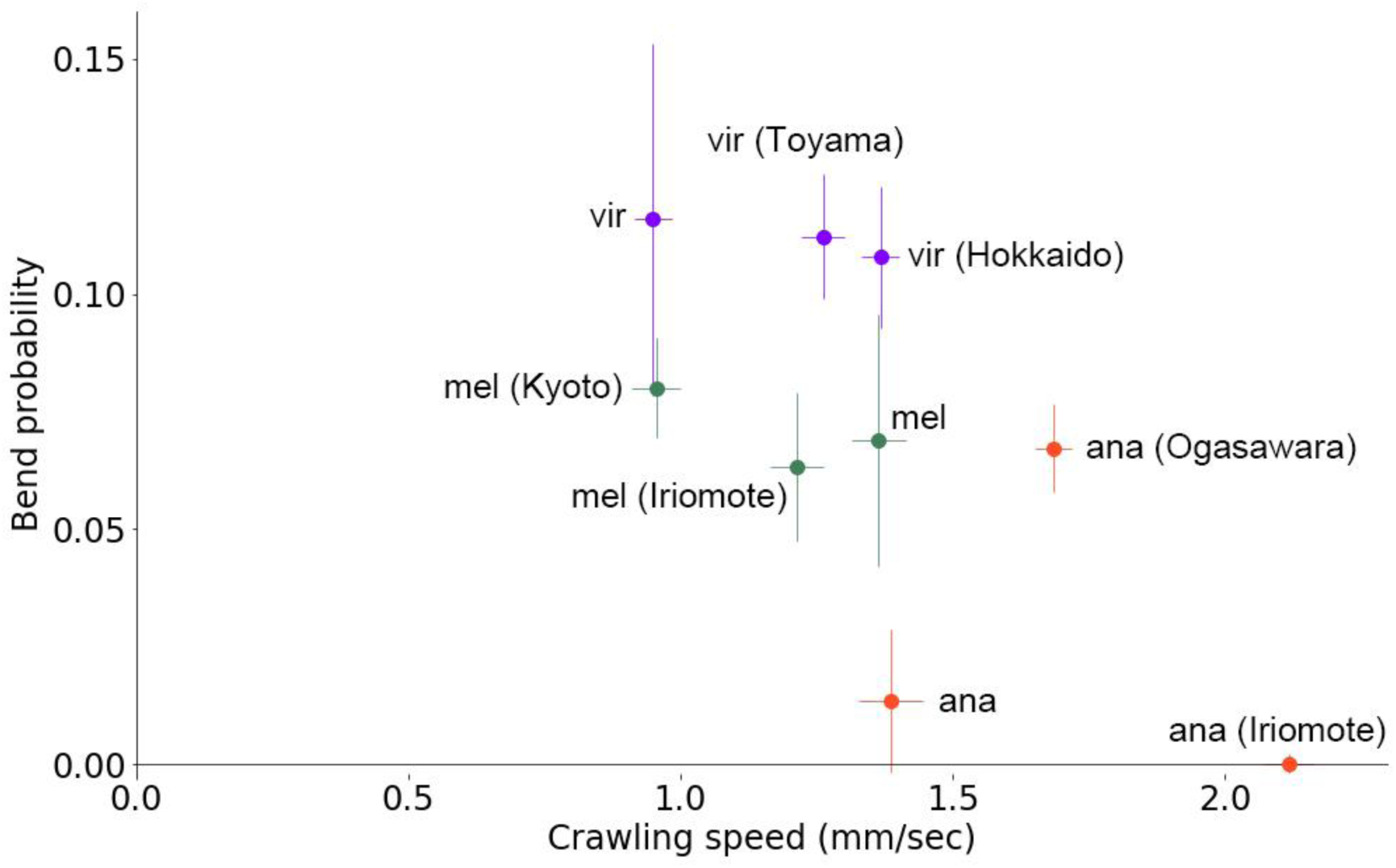
Intraspecific comparison in larval locomotion. Scatter plot of bend probability at 24°C against crawling speed at 24°C of nine strains from three species. Sample numbers are the following: *Dvir*: n=44; *Dvir* (Hokkaido): n=32; *Dvir* (Toyama): n=35; *Dmel*: n=33; *Dmel (Kyoto)*: n=47; *Dmel (Iriomote)*: n=33; *Dana*: n=33; *Dana (Ogasawara)*: n=40; *Dana (Iriomote)*: n=20.

### Crawling distance is related to crawling speed and bend probability

In the interspecies variability, there is a negative correlation between the crawling speed and bend probability (Figure 3E). We noticed that this correlation could reflect the control of the crawling distance by a coordinated change of the crawling speed along with the bend probability. The bend probability is negatively correlated with the crawling distance since frequent bending shortens the distance larvae progress in one direction (Figure 5A) whereas it is obvious that the crawling speed is positively correlated with the crawling distance (Figure 5B). Accordingly, high crawling speed and low bend probability, which are a combination appeared in the negative correlation of them (Figure 3E) both contribute to the increase in the crawling distance. To check whether these geometrical speculations hold in the locomotion of larvae, we examined the relationship between the kinematic parameters and the crawling distance (Figure 5C and 5D). Consistent with the conjecture, the crawling distance is larger when either the bend probability is lower (Figure 5C; Pearson correlation = -0.78) or the crawling speed is higher (Figure 5D; Pearson correlation = 0.67). These observations imply that the coordinated changes in the crawling speed and bend probability would be related to the change in the crawling distance of larvae of each species.

**Figure 5.**
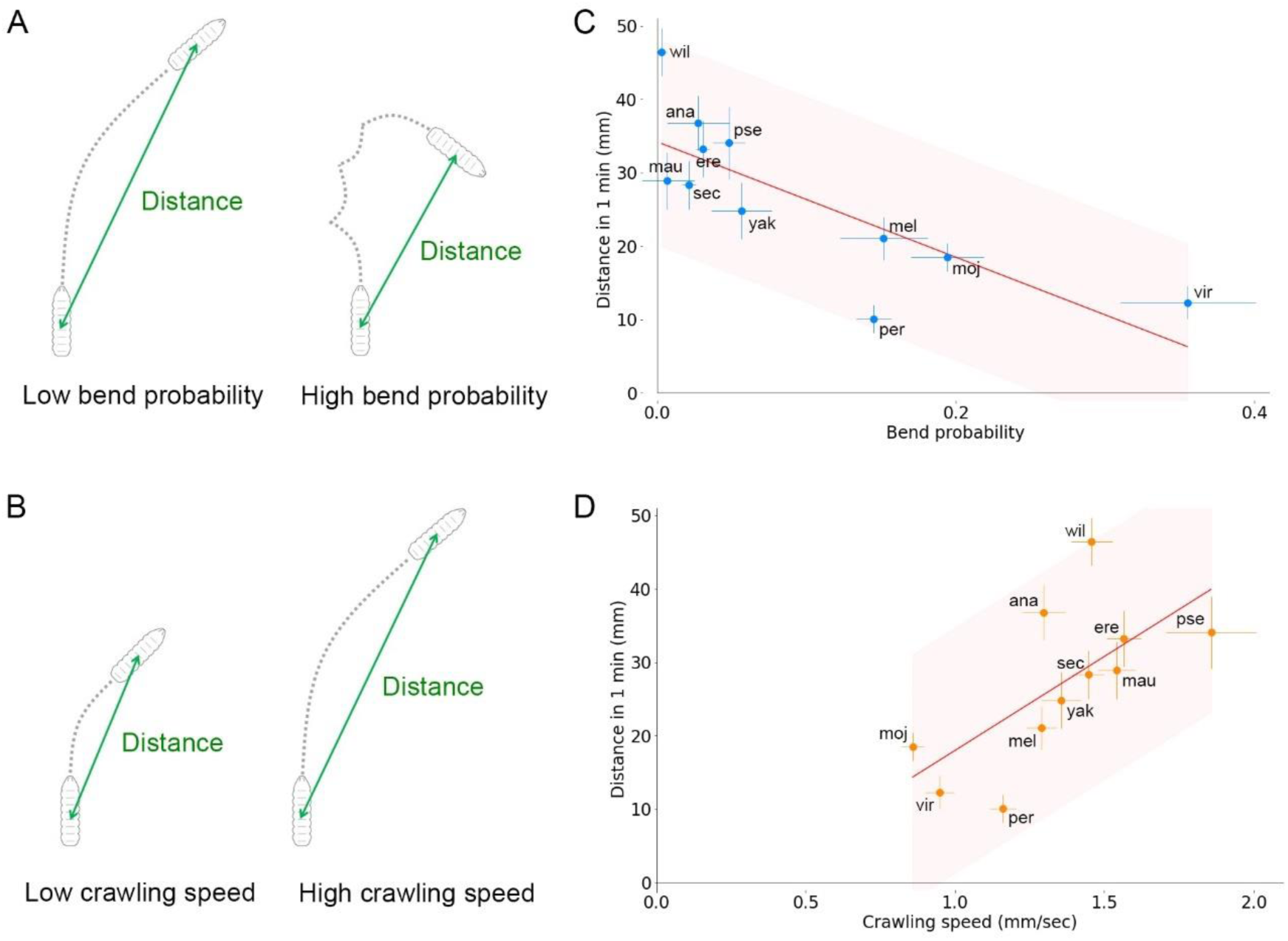
Relationship between crawling distance and the two kinematic parameters in the 11 Drosophila species. (A) Schematics of the relationship between the bend probability and crawling distance. (B) Schematics of the relationship between the crawling speed and crawling distance. (C) Scatter plot of the bend probability and crawling distance in one minute in the 11 species. (D) Scatter plot of the crawling speed and crawling distance in one minute in the 11 species. The source locomotion data are the same as in Figure 3C and 3D. In C and D, the red lines show the linear regression functions and the shaded areas represent the 95% confidence bands. The point estimates of the Pearson correlation and their 95% confidence intervals are -0.78 and [-0.94, -0.34] in C and 0.67 and [0.11, 0.91] in D.

### No relationship between the kinematics of larval locomotion and the body length nor phylogenetic relationship

Next, we tried to find factors that relate to the diversification of larval kinematics among species. Among the factors in the morphological differences and the ecological diversity that might be involved in the kinematics diversity (Theunissen et al., 2015) we examined the body length of larvae as a morphological factor and habitat temperature as an ecological factor. A previous study reported an allometric relationship between the body size and the crawling speed in *Diptera* larvae (Berrigan & Pepin, 1995). The authors analyzed larvae in the order *Diptera*, the average size of which spans from 3.7 mm (*Dmel*) to 15.9 mm (*Sarcophaga bullata*). We tested whether the relationship between the body length and the crawling speed also holds within the genus *Drosophila*, a subgroup of the order *Diptera*. The length of the *Drosophila* larvae we used spanned from 3.49 ± 0.05 mm (*Dyak*) to 5.68 ± 0.16 mm (*Dpse*) (Supplementary Figure 2A). We found no significant relationship between the body length and crawling speed (Supplementary Figure 2B; Pearson correlation = 0.04, p = 0.91) and between the body length and the bend probability (Supplementary Figure 2C; Pearson correlation = 0.55, p = 0.077). These data suggest that larval length is not a significant factor for larval kinematics variation within the genus *Drosophila*.

Phylogenetic relationship can also be a factor that affects the kinematics. To test this issue, we focused on two species groups: the obscura species group and replete species group. *Dper* and *Dpse* belongs to the obscura species group (See Figure 11A). The kinematics of these two sister species are separated in the distribution of the crawling speed and bend probability (Figure 3E). Similarly, two species of the replete species group, *Dvir* and *Dmor* exhibit distinct kinematics (Figure 3E). These observations suggest that phylogenetic relationship is not a major factor in the divergence of the kinematics.

### Relationship between the kinematics of larval locomotion and habitat temperature of the *Drosophila* species

Habitat temperature is one of the critical factors influencing species traits (Schmidt-Nielsen, 1997). To test the possible roles of habitat temperature in the evolution of *Drosophila* larval locomotion, we examined the relationship between habitat temperatures and the locomotion kinematics in the genus *Drosophila*. The habitat regions of the 11 species were obtained from the literature (Brake & Baechli, 2008; Makino & Kawata, 2012; Markow & O’Grady, 2005), and climate temperature data were obtained from a global climate dataset, WORLDCLIM (Hijmans et al., 2005) (Figure 6A, Supplementary Figure 3). We examined the relation between larval kinematics (bend probability or crawling speed) and indices of habitat temperatures (Figure 6B and 6C). We used the mode (the most frequent value) of three indices of habitat temperatures: average, maximum, and minimum temperature. In the minimum habitat temperature data of *Dvir* and *Dpse*, we noticed that the mode temperatures were below 0°C, which cannot be a representative habitat temperature for them (Supplementary Figure 3). Accordingly, for a milder and more representative index, we used the warmest peaks in the minimum habitat temperature histogram (“Tmin” in Supplementary Figure 3). Considering uneven distribution of flies within the region demarcated in Supplementary Figure 3, the temperature of the warmest peak in the minimum habitat temperature may represent the coldest temperature the majority of the population of each species experiences over many years.

**Figure 6.**
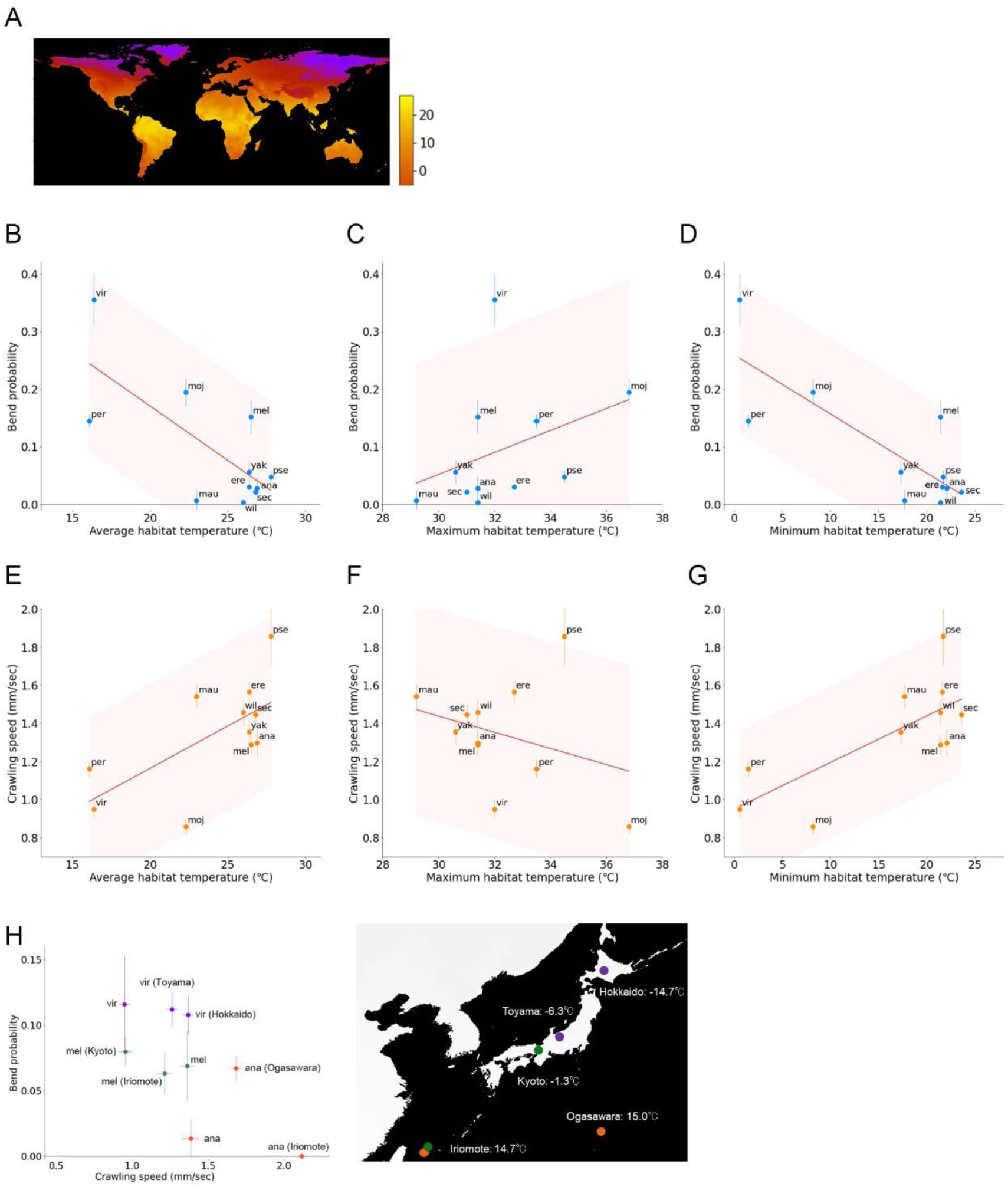
Relationship between the kinematics of larval locomotion and habitat temperature of the Drosophila species. (A) A world map of minimum habitat temperature. (B - D) Scatter plot of the bend probability of the 11 species at 24°C against the average (B), maximum (C), and minimum (D) habitat temperatures. (E - G) Scatter plot of the crawling speed of the 11 species at 24°C against the average (E), maximum (F), and minimum (G) habitat temperatures. In B to G, median ± sem are shown and the source locomotion data are the same as in Figure 3C and 3D. The red lines show the linear regression functions and the shaded areas represent the 95% confidence bands. The point estimates of the Pearson correlation and their 95% confidence intervals are -0.73 and [-0.93, -0.23] in B, 0.37 and [-0.30, 0.79] in C, and -0.81 and [-0.95, -0.41] in D, 0.66 and [0.09, 0.90] in E, -0.31 and [-0.77, 0.36] in F, and 0.74 and [0.25, 0.93] in G. (H) Left: A scatter plot of intraspecific comparison (the same as Figure 4). Right: A map of Japan representing the minimum habitat temperatures of the six strains shown in the left panel

We found that the bend probability and crawling speed are both correlated with average habitat temperature Tave (Figure 6B and 6E. Pearson correlation: bend probability vs Tave = -0.73, crawling speed vs Tave = 0.66). We further analyzed whether the maximum and minimum temperature contribute to the correlation. Maximum habitat temperature, Tmax shows no obvious correlation to the kinematics (Figure 6C and 6F. Pearson correlation: bend probability vs Tmax = 0.37, crawling speed vs Tmax = -0.31). In contrast, minimum habitat temperature, Tmin, exhibit stronger correlation than Tmax (Figure 6D and 6G. Pearson correlation: bend probability vs Tmin = -0.81, crawling speed vs Tmin = 0.74). Even after omitting extreme values (*Dvir* in the bend probability data and *Dpse* in the crawling speed data), the correlation remained high (Pearson correlation: the bend probability vs Tmin (without *Dvir*) = -0.69, the crawling speed vs Tmin (without *Dpse*) = 0.78). Furthermore, even considering multiple comparisons (four factors: larval length, Tave, Tmax and Tmin), the correlations between the kinematic parameters and Tmin were statistically significant (p = 0.039 in crawling speed vs Tmin, p = 0.0010 in bend probability vs Tmin; Bonferroni correction).

The range (or variability) of habitat temperature could also be a critical factor to determine larval kinematics because species that inhabits in highly variable temperature area would be insensitive to the change in the ambient temperature whereas those breeding in a narrow range of temperature would be sensitive to the small shift in the ambient temperature. To test this point, we examined the relationship between the larval kinematics and the range of habitat temperature, the difference between maximum and minimum temperature in their habitat area (Supplementary Figure 4). The range of habitat temperature is correlated with both the bend probability (Pearson correlation = 0.65, p = 0.030) and the crawling speed (Pearson correlation = -0.69, p = 0.0019). Consequently, the range of habitat temperature could affect the larval kinematics. In the following analysis, we will focus on the minimum temperature Tmin since it showed the most evident correlation to the larval kinematics.

It should be noted that the strains we used have been kept at the housed stocks at 23 or 20°C (See Materials and Methods), which might affect the innate behaviour in wild type strains. However, comparison between time period after speciation (at least million years ∼ 10^7^ generations (Campbell & Ganetzky, 2012; Russo et al., 1995) and that in laboratories (100 years ∼ 10^3.5^ generations) indicates that the duration in laboratories occupies as small as 0.03% of the time for the evolution of these *Drosophila* strains. In addition, the rearing temperatures in the stock centres were within the range of habitat temperature of each species (Supplementary Figure 3). These rearing conditions imply that the strains we collected from the stock center should possess innate behaviour that was evolved in their habitat.

To sum, larvae of species inhabiting moderate environments (where Tmin is 15 to 25°C) showed low bend probability and fast crawling, or long crawling distance, whereas those inhabiting cold environments (where Tmin is 0 to 10°C) exhibited frequent bending and slow crawling, or short crawling distance. Especially, species that has lower Tmin than the ambient temperature in this assay (24°C) showed shorter crawling distance, which implies that excess heat stimuli to these species should reduce their crawling distance.

In the analysis above, we estimated the indices of habitat temperature from the temperature data in the entire potential habitat of each species because the original habitat of each strain was unclear. To further check the correlation between the larval kinematics and habitat temperature, we examined the relationship between the kinematics and habitat temperature among the strains of which habitat were recorded (Figure 4 and 6H). We used two *Dmel* strains (collected at Kyoto and Iriomote in Japan), two *Dvir* strains (collected at Hokkaido and Toyama in Japan), and two *Dana* strains (collected at Ogasawara and Iriomote in Japan). The habitat temperature at these location for collecting flies is diverse (Figure 6H right). Intriguingly, the strains originated from the north part of Japan, where the minimum habitat temperature is low, exhibit slower crawling and high bend probability whereas those from the south part of Japan, where the minimum habitat temperature is high, show the opposite trend (Figure 6H left), which is consistent with the observation based on the temperature of the worldwide statistics (Figure 6D and 6G). Consequently, these observations imply that habitat temperature should be one of the leading factors in sculpting the kinematics of larval locomotion in the genus *Drosophila*.

### Correlation between bend probability and habitat temperature among *Drosophila* species at high ambient temperature

So far, we have analyzed larval locomotion at 24°C, which is close to the rearing temperature (See Materials and Methods). Larvae of *Dmel* are known to crawl faster at 32°C than at 25°C (Ohyama et al., 2013). So, we next examined whether the relation between the kinematics indices and habitat temperature holds or not, and how the kinematics indices are changed, at a higher ambient temperature. We performed the same set of measurements and analysis of larval locomotion at 32°C (Figure 7 and 8). The interspecies comparison showed that the bend probabilities at 32°C were correlated with minimum habitat temperature (Figure 7B; Pearson correlation = -0.85, p = 0.0009). To examine the effects of ambient temperature on the bend probability in detail, we compared the differences of bend probability at 24°C and 32°C among the species (Figure 7C). We found no significant relationship between the change in bend probability and the minimum habitat temperature (Figure 7C; Pearson correlation = 0.40, p = 0.22). Seven species showed no significant changes in the bend probability at 32°C (Mann-Whitney U-test of *Dmoj*, *Dper*, *Dpse*, *Dsec*, *Dmau*, *Dana* and *Dyak* in Figure 7A - 7D), while *Dvir* and *Dmel* showed a decrease in the bend probability at 32°C, and *Dwil* and *Dere* exhibited an increase (Mann-Whitney U-test in Figure 7A - 7C, 7E and 7F). Accordingly, we concluded that while the shift of the bend probability between the distinct temperatures is diverse among the *Drosophila* species, the overall trend between the bend probability in relation to habitat temperature holds at the higher ambient temperature of 32°C.

**Figure 7.**
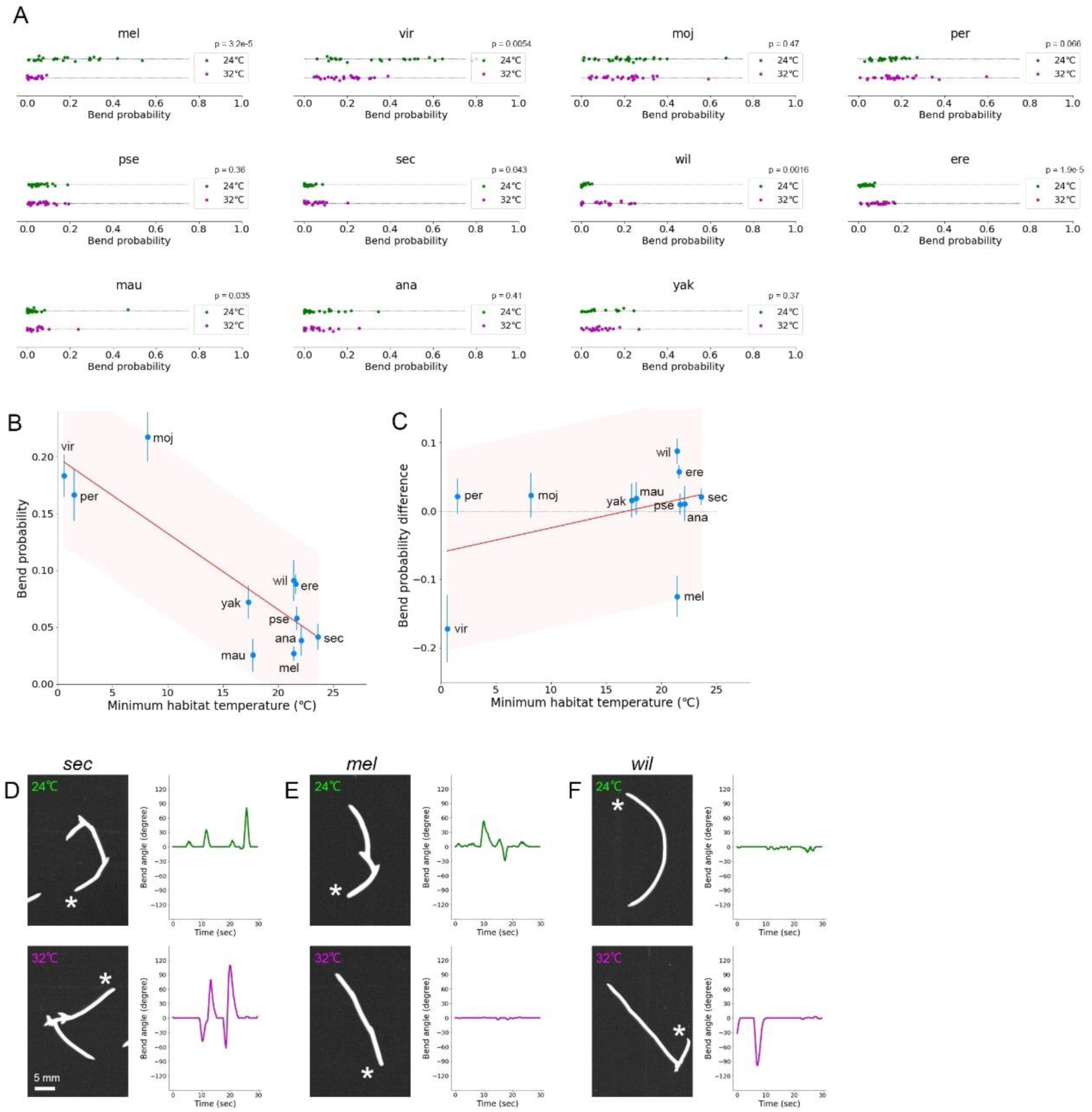
Bend probability in larval locomotion in the 11 *Drosophila* species at 32°C. (A) Bend probability of the 11 species at 24°C and 32°C. The data at 24°C are the same as in Figure 3C. Sample numbers at 32°C are the following: *Dvir*: n=25; *Dmel*: n=20; *Dmoj*: n=29; *Dper*: n=27; *Dpse*: n=25; *Dsec*: n=20; *Dwil*: n=21; *Dere*: n=24; *Dmau*: n=17; *Dana*: n=23; *Dyak*: n=19. P values present the results of Mann-Whitney U-test. (B) Scatter plot of bend probability at 32°C against the minimum habitat temperature, Tmin. (C) Scatter plot of the difference in bend probability between 32°C and at 24°C (bend probability at 32°C – bend probability at 24°C) against the minimum habitat temperature Tmin. In B and C, median ± sem is shown. The red lines show the linear regression functions and the shaded areas represent the 95% confidence bands. The point estimates of the Pearson correlation and their 95% confidence intervals are -0.850 and [-0.96, -0.51] in B and 0.40 and [-0.26, 0.81] in C. (D-F) Examples of trajectories (left) and the bend angle (right) at 24°C (top) and 32°C (bottom) of larval locomotion of each species (D: *Dsec*, E: *Dmel*, and F: *Dwil*).

**Figure 8.**
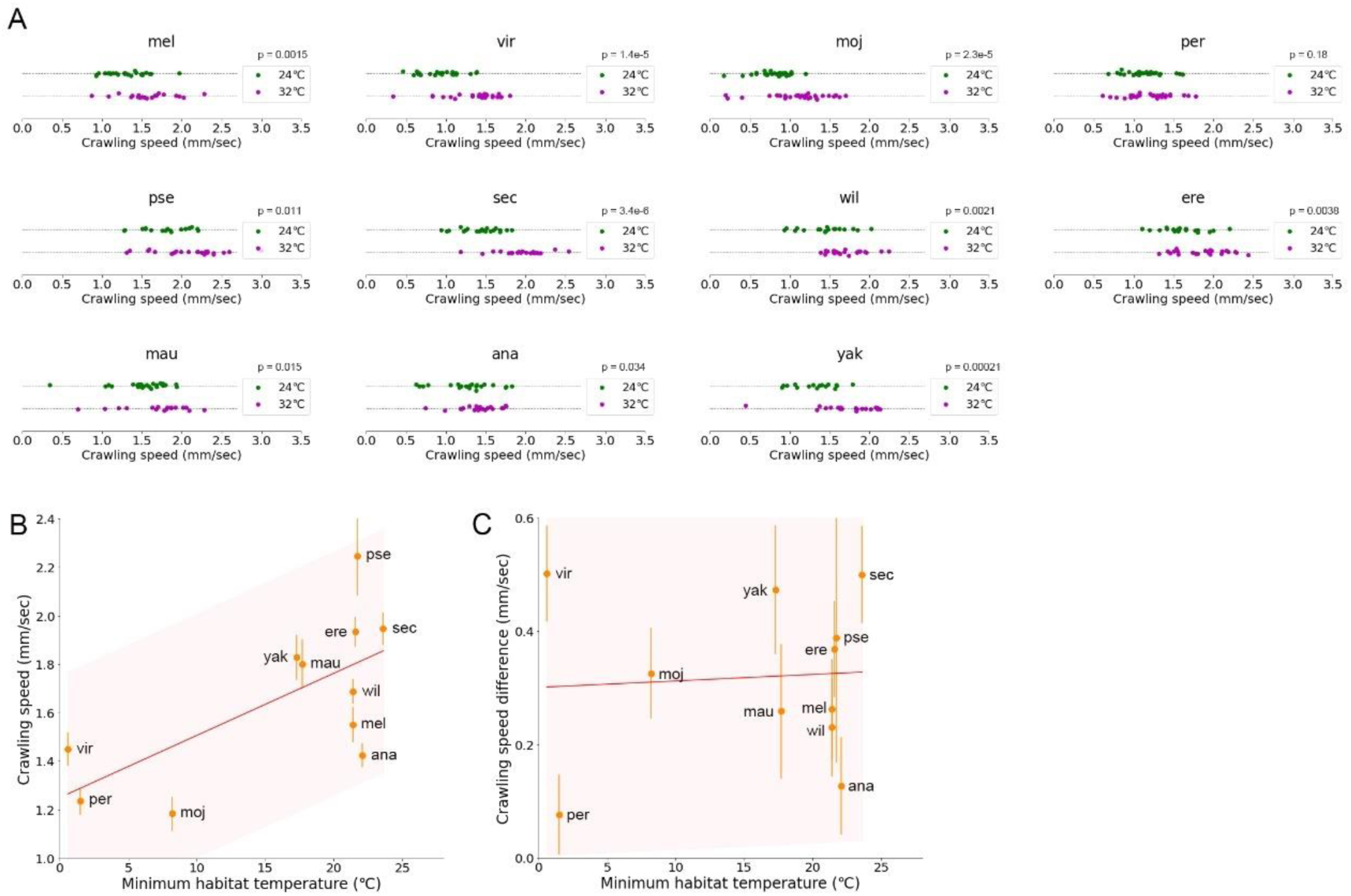
Crawling speed of larval locomotion in the 11 *Drosophila* species at 32°C. (A) Crawling speed of the 11 species at 24°C and 32°C. The data at 24°C are the same as in Figure 3D. Sample numbers at 32°C are the following: *Dvir*: n=25; *Dmel*: n=20; *Dmoj*: n=29; *Dper*: n=27; *Dpse*: n=25; *Dsec*: n=20; *Dwil*: n=21; *Dere*: n=24; *Dmau*: n=17; *Dana*: n=23; *Dyak*: n=19. P values present the results of Mann-Whitney U-test. (B) Scatter plot of the crawling speed at 32°C against the minimum habitat temperature Tmin. (C) Scatter plot of the difference of the crawling speed between at 32°C and at 24°C (crawling speed at 32°C – crawling speed at 24°C) against the minimum habitat temperature Tmin. In B and C, median ± sem is shown. The red lines show the linear regression functions and the shaded areas represent the 95% confidence bands. The point estimates of the Pearson correlation and their 95% confidence intervals are 0.67 and [0.12, 0.91] in B and 0.07 and [-0.56, 0.64] in C.

### Correlation between crawling speed and habitat temperature among *Drosophila* species at high temperature

Next, we analyzed the crawling speed at 32°C (Figure 8A). Similar to the bend probability, we found that the crawling speed at 32°C was correlated with the minimum habitat temperature among the species, as it was at 24°C (Figure 8B; Pearson correlation = 0.67, p = 0.024). To examine the effects of ambient temperature on the crawling speed in detail, we compared the difference in the crawling speed at 24°C to 32°C among the species (Figure 8C). In contrast to the case in the bend probability, shifts in the crawling speed among the species were common: ten species (all the species but *Dper*) showed a significant increase in crawling speed at 32°C (Mann-Whitney U-test in Figure 8A). *Dper* also exhibited an increase in the crawling speed but it was not statistically significant (Mann-Whitney U-test in Figure 8A). Accordingly, at 32°C, larvae of all the species we tested increased their speed of crawling, which might reflect a general demand to avoid malfunctions in metabolic reactions and dehydration of the body. We also tested the relationship between the change in the speed at 32°C and the minimum habitat temperature for each species (Figure 8C). We found no significant correlation between the speed change and the habitat temperature (Figure 8C; Pearson correlation = 0.07, p = 0.84), which also implies that the speeding-up at high temperature is a general requirement among the species for the larval survival. Accordingly, whereas the details of shifts are different (diverse shift in the bend probability and common shift in the crawling speed among the species), the overall relationship between the kinematic indices and minimum habitat temperature also holds at the higher ambient temperature of 32°C.

### A kinematic trend of larval locomotion in the genus Drosophila

To see any gross trend in the shift of larval kinematics in the different species at distinct ambient temperatures, we plotted the kinematics of individual larvae of all the species (Figure 9). At the ambient temperature of 24°C, a trend of kinematics was observed in which the data points were distributed from low crawling speed / high bend probability (the top-left corner in the plot) to high crawling speed / low bend probability (the bottom-right corner in the plot) as the minimum habitat temperature increased (Figure 9A). Intriguingly, the kinematics indices between distinct species were not segregated but rather overlapped. This continuum property can also be observed in the kinematics data at the ambient temperature of 32°C (Figure 9B). According to the modern phylogenetic concept (Markow & O’Grady, 2005), the 11 genus *Drosophila* species in our study consisted of nine subgenus *Sophophora* species (all but *Dvir* and *Dmoj*) and two *non-Sophophora* species (*Dvir* and *Dmoj*). In our plots of kinematics, species in the *non-Sophophora* (*Dvir* and *Dmoj*) were located at the left side in the continuum (Figure 9). This observation implies that the kinematics indices of larvae in *Sophophora* species in the genus *Drosophila* took values in this continuum and kinematics indices in *non-Sophophora* species in the genus *Drosophila* have diverged along this continuum by adaptation to habitat temperature during evolution.

**Figure 9.**
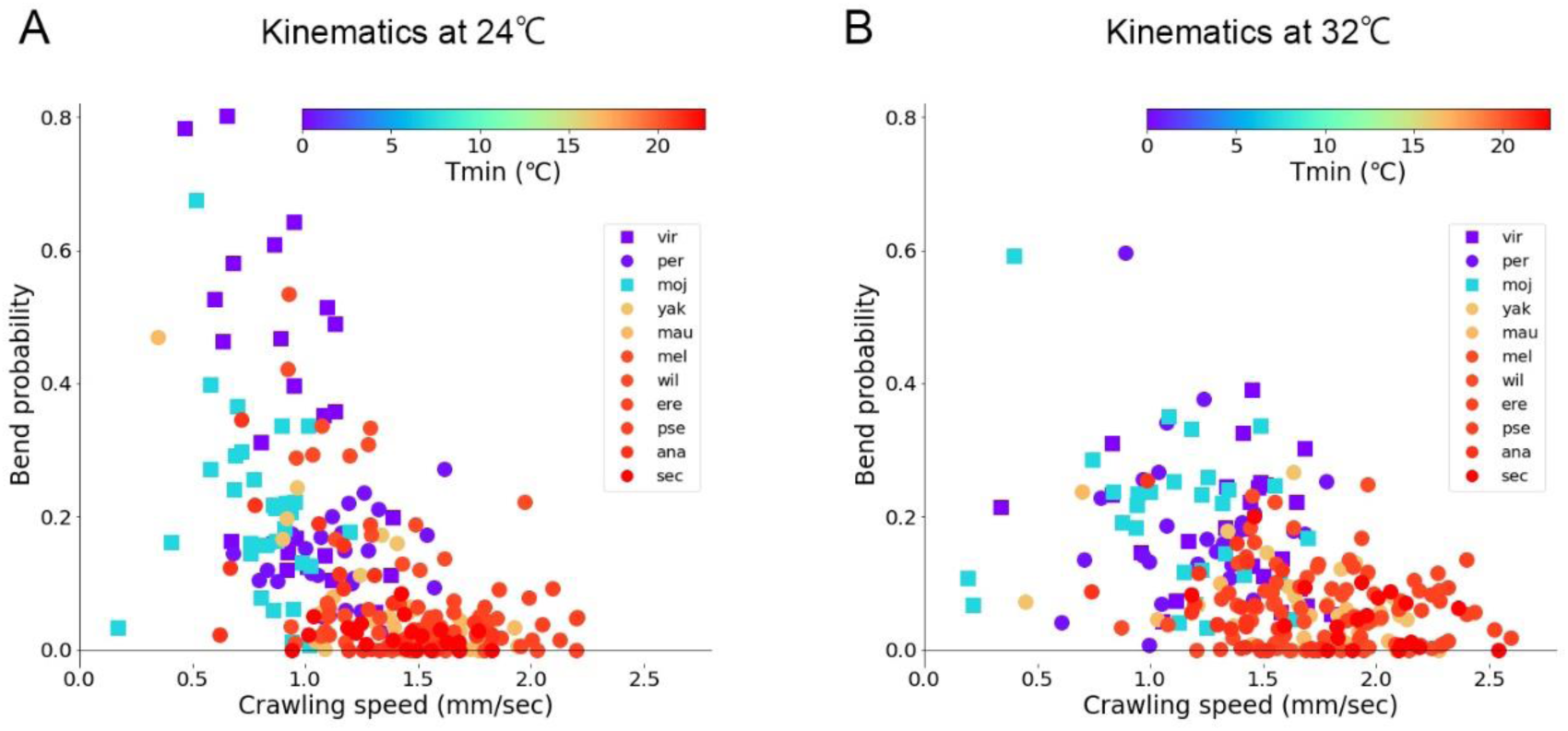
Scatter plot of kinematics in the 11 species at 24°C and 32°C. (A and B) Two dimensional plots of the crawling speed and bend probability. Each point corresponds to data of a single larva of the species. The colour of markers denotes minimum habitat temperature (Tmin) as shown in the color bar. Circles denote *Sophophora* species and squares denote *non-Sophophora* species. The source locomotion data are the same as Figure 3, 7, and 8. A: 24°C, B: 32°C.

### No correlation between crawling speed and habitat temperature among *Drosophila* species at an extreme ambient temperature of 40°C

Temperatures of 24°C and 32°C are within the range of natural habitat temperatures of most of the species we tested (Supplementary Figure 3). We next examined larval crawling at an extreme temperature and investigated the relation between the kinematics indices and minimum habitat temperature. Too high a temperature can be noxious for many animals, including *Drosophila* larvae. When *D. melanogaster* larvae are stimulated with a probe heated to 42°C, they exhibit a stereotyped rolling motion as an escape behaviour (Dason et al., 2019; Petersen et al., 2018; Tracey et al., 2003). At below 40°C, on the other hand, larvae do not exhibit rolling behaviour (Petersen et al., 2018), which allows us to study crawling kinematics in this semi-noxious extreme environment. We recorded larval locomotion of the *Drosophila* species at the ambient temperature of 40°C, which corresponds to the highest edge of maximum habitat temperature histograms for many of the species (Supplementary Figure 3). We measured the bend probability and crawling speed of the 11 *Drosophila* species (Supplementary Figure 5 and 6). At the ambient temperature of 40°C, neither the bend probability nor crawling speed was correlated with minimum habitat temperature (Figure 10A and 10B; Pearson correlation: bend probability at 40°C vs Tmin = 0.17, p = 0.61; crawling speed at 40°C vs Tmin = -0.17, p = 0.61). Since 40°C is close to the maximum habitat temperature rather than the minimum habitat temperature, we analyzed the relation between the kinematics indices and maximum habitat temperature instead of minimum habitat temperature. However, neither the bend probability nor crawling speed was correlated with maximum habitat temperature (Pearson correlation: bend probability at 40°C vs maximum habitat temperature = -0.13, p = 0.71; crawling speed vs maximum habitat temperature = -0.10, p = 0.77). Accordingly, the relation between the kinematic indices and habitat temperature does not hold at the extreme ambient temperature of 40°C. This observation suggests that the influence of habitat temperature on the evolution of the locomotion kinematics is restricted within a specific range of ambient temperatures. To examine the effects of the extreme ambient temperature on the kinematics indices in detail, we examined the shift in the bend probability at the distinct temperatures of 40°C and 32°C among the species (Supplementary Figure 5). In seven species, bend probability increased at 40°C. Three species that inhabit moderate temperature areas (*Dvir*, *Dper* and *Dmoj*) and one species that lives on an isolated island (*Dsec*) showed little change in the bend probability, which may reflect a distinct strategy in evolution to cope with the semi-noxious environment. We also examined the shift in the crawling speed at the distinct temperatures of 40°C and 32°C among the species (Supplementary Figure 5). We found the crawling speed was reduced at 40°C in all the species (Supplementary Figure 6), which might be a common adaptation of the *Drosophila* larval locomotion at the extreme ambient temperature and/or due to an abnormal physiological reaction at the semi-noxious temperature. The similar tendency, the increase in the bend probability and the decrease in the crawling speed at 40°C, was also observed when compared with the kinematics at 24°C (Figure 10C and 10D). Intriguingly, in some species, backward crawling, which seldom occurs at 32°C, can be observed at 40°C (Figure 10E). Consequently, these observations show that at the extreme ambient temperature of 40°C, the relation between the kinematics indices and habitat temperature does not hold and larvae exhibit common (increase in the crawling speed) and diverse (changes in the bend probability and generation of backward crawling) shifts in the locomotion kinematics among the *Drosophila* species.

**Figure 10.**
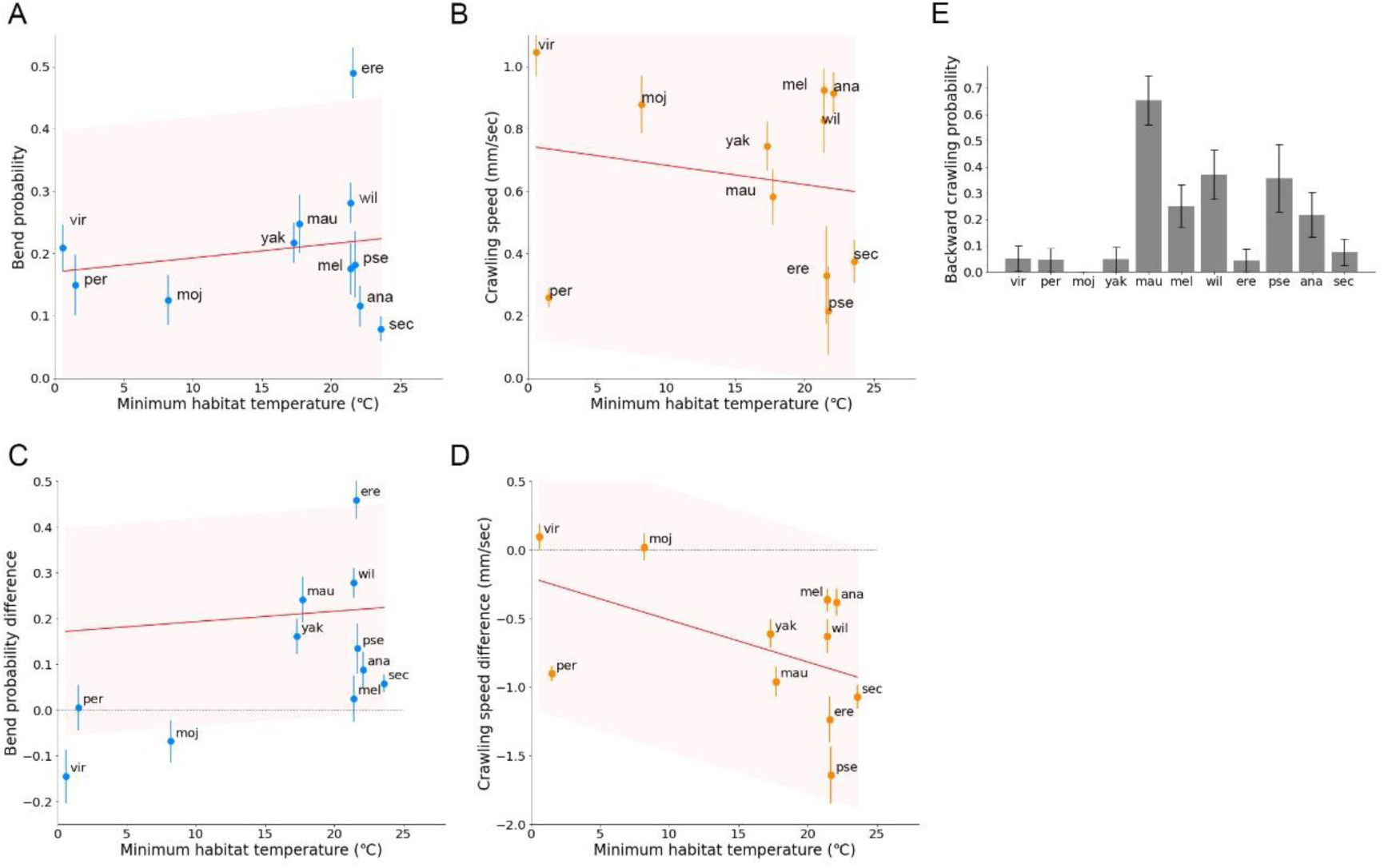
Larval behaviour in the 11 species at 40°C. (A) Scatter plot of the bend probability at 40°C against the minimum habitat temperature Tmin. (B) Scatter plot of the crawling speed at 40°C against the minimum habitat temperature Tmin. (C) Scatter plot of the difference of the bend probability between at 40°C and at 24°C (bend probability at 40°C – bend probability at 24°C) against the minimum habitat temperature Tmin. (D) Scatter plot of the difference of the crawling speed between at 40°C and at 24°C (crawling speed at 40°C – crawling speed at 24°C) against the minimum habitat temperature Tmin. Median ± sem is shown in A - D. Sample numbers for data at 40°C are the following: *Dvir*: n=20; *Dmel*: n=28; *Dmoj*: n=23; *Dper*: n=21; *Dpse*: n=16; *Dsec*: n=27; *Dwil*: n=27; *Dere*: n=22; *Dmau*: n=26; *Dana*: n=23; *Dyak*: n=20. Sample numbers for data at 24°C are the same as Figure 3C and 3D. The red lines show the linear regression functions and the shaded areas represent the 95% confidence bands. The point estimates of the Pearson correlation and their 95% confidence intervals are 0.17 and [-0.48, 0.70] in A and -0.17, [-0.70, 0.48] in B, 0.17, [-0.48, 0.70] in C, and -0.50, [-0.84, 0.15] in D. (E) Backward crawling probability at 40°C of each species. Probability ± standard error based on the binomial distribution are shown. Sample numbers in E are the following: *Dvir*: n=20; *Dmel*: n=28; *Dmoj*: n=23; *Dper*: n=22; *Dpse*: n=14; *Dsec*: n=27; *Dwil*: n=27; *Dere*: n=23; *Dmau*: n=26; *Dana*: n=23; *Dyak*: n=21.

### Phylogenetic analyses of interspecies differences in kinematics

Finally, to obtain a hint on how the kinematics parameters diversified across the species have developed during the evolutionary history of the genus *Drosophila*, we conducted Bayesian phylogenetic analyses using RevBayes v. 1.0.7 (Hohna et al., 2016; Höhna et al., 2017). At first, we inferred phylogenetic trees of the 11 *Drosophila* species based on eight nuclear genes used previously (Turelli et al., 2018) (See Methods for detail). Then by using our kinematics dataset, we estimated the following three factors in the phylogenetic tree by Bayesian inference: the rate of evolution at each branch of the phylogenetic trees, relative rates of evolution among the kinematic parameters, and the correlation between the kinematic parameter evolutions (Kalay et al., 2020). To perform the inference, we constructed a data matrix of eight parameters (the bend probability at 24, 32 and 40°C, the crawling speed at 24, 32 and 40°C, backward crawling probability at 40°C and the body length), for each of the 11 *Drosophila* species. We assumed that these parameters evolve under a multivariate Brownian-motion model (Huelsenbeck & Bruce, 2003; Kalay et al., 2020; Lartillot & Poujol, 2011) (See Methods for detail). We estimated the three factors (the evolution rates at phylogenetic trees, relative evolution rates among the kinematic parameters, and the correlation between the parameter evolutions) by running a Markov Chain Monte Carlo simulation.

The phylogenetic analyses indicated that the rates of evolution in the kinematics are highly diverse over branches in the phylogenetic tree (Figure 11A and Supplementary Figure 7). In addition, the evolutionary rates are distinct among the eight kinematics parameters (Figure 11B). The bend probability at 24°C and the probability of backward crawling at 40°C have a relatively high evolution rate, which might reflect large diversification of these two parameters among the species (Figure 9A and 10E).

Some kinematics traits might have evolved cooperatively. To test this possibility, we calculated the correlation of the evolutionary changes between the kinematic parameters. In the correlation analyses of all pairs of the parameters, the crawling speed at 24°C and the crawling speed at 32°C were the most correlated (Figure 11C). On the other hand, the evolutions of the bend probability between at 24°C and 32°C were less correlated. This observation is consistent with the notion that while changes in the bend probability from 24°C to 32°C among the species are diverse (Figure 7C), all the species show an increase in the crawling speed by the temperature shift (Figure 8C). Consequently, the phylogenetic analyses imply that the kinematics indices of larval locomotion have evolved differently in distinct branches of a phylogenetic tree with keeping a correlation between specific locomotion traits such as crawling speed at distinct temperatures.

## Discussion

In this work, we investigated interspecies differences in larval locomotion in the genus *Drosophila*. We used the bend probability and crawling speed as measures to examine larval locomotion. Despite the similar appearance of larval bodies in the different species, the kinematics of larval locomotion is diverged (Figure 2). The body length is not a leading factor for the diversity of kinematics (Supplementary Figure 2). Phylogenetic relationship is also not a major determinant for the kinematics (Figure 3E and 11A). Considering a previous study showing that phylogenetic relationship does not correlate with the divergence in the morphology of larval neuromuscular junctions (Campbell & Ganetzky, 2012), genetic drift with random accumulation of neutral mutations is unlikely to underlie the divergence in the larval crawling patterns. In contrast, habitat temperature correlates with both the bend probability and crawling speed at both 24°C and 32°C ambient temperature (Figure 12), which implies the kinematics indices are adapted to ambient temperature in evolution. Phylogenetic analysis by Bayesian inference suggests that the rates of evolution are divergent among the branches of the phylogenetic tree (Figure 11A). Among the kinematic parameters, the bend probability at the ambient temperature of 24°C and the backward crawling probability at 40°C have relatively higher evolution rates than the others and the evolution of the crawling speed at 24°C and 32°C are correlated (Figure 11B and 11C). Regarding the questions raised in the Introduction section, our results suggest the following: (1) Locomotion kinematics of larvae is divergent among *Drosophila* species. (2) The habitat temperature is more related to the kinematics indices (the bend probability and crawling speed) than the body length (Figure 12).

**Figure 11.**
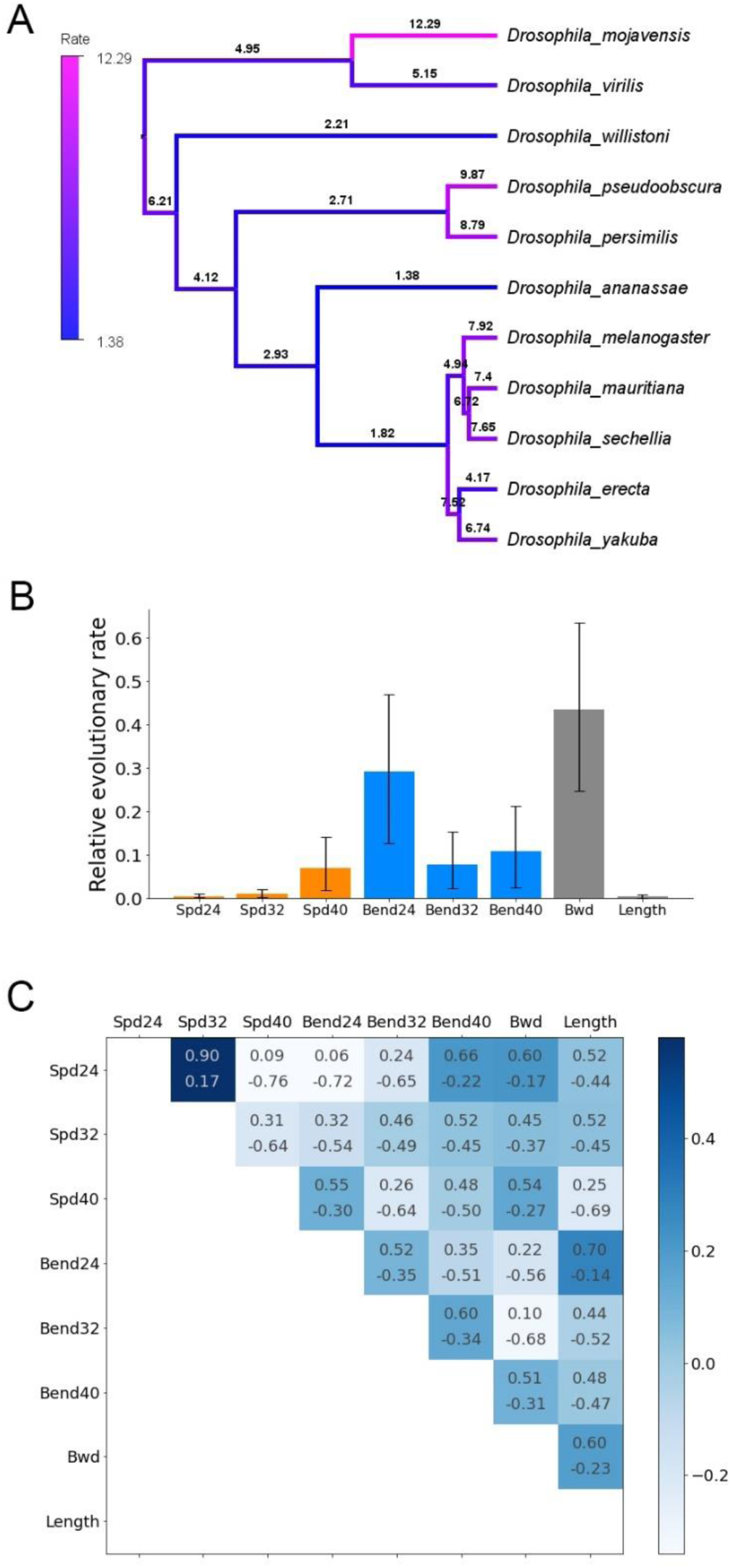
Phylogenetic analyses of larval locomotion in the genus *Drosophila*. (A) Phylogenetic tree estimated from Bayesian inference. The topology is inferred by coding sequences of eight genes of the 11 *Drosophila* species. As a prior distribution for the Bayesian inference, an uncorrelated exponential (UCED) relaxed-clock model was used. Colour and a value of each branch denote the relative rate of the evolution of the eight kinematic traits. (B) Relative evolution rates of the eight kinematics traits estimated by the Bayesian inference. Mean ± 95% highest posterior density is shown. (C) Correlation between the evolution of eight traits. Numbers represent the range of 95% highest posterior density. Abbreviations in (B) and (C) are the following: Spd24: crawling speed at 24°C, Spd32: crawling speed at 32°C, Spd40: crawling speed at 40°C, Bend24: bend probability at 24°C, Bend32: bend probability at 32°C, Bend40: bend probability at 40°C, Bwd: probability of backward crawling, Length: axial body length of larvae.

**Figure 12.**
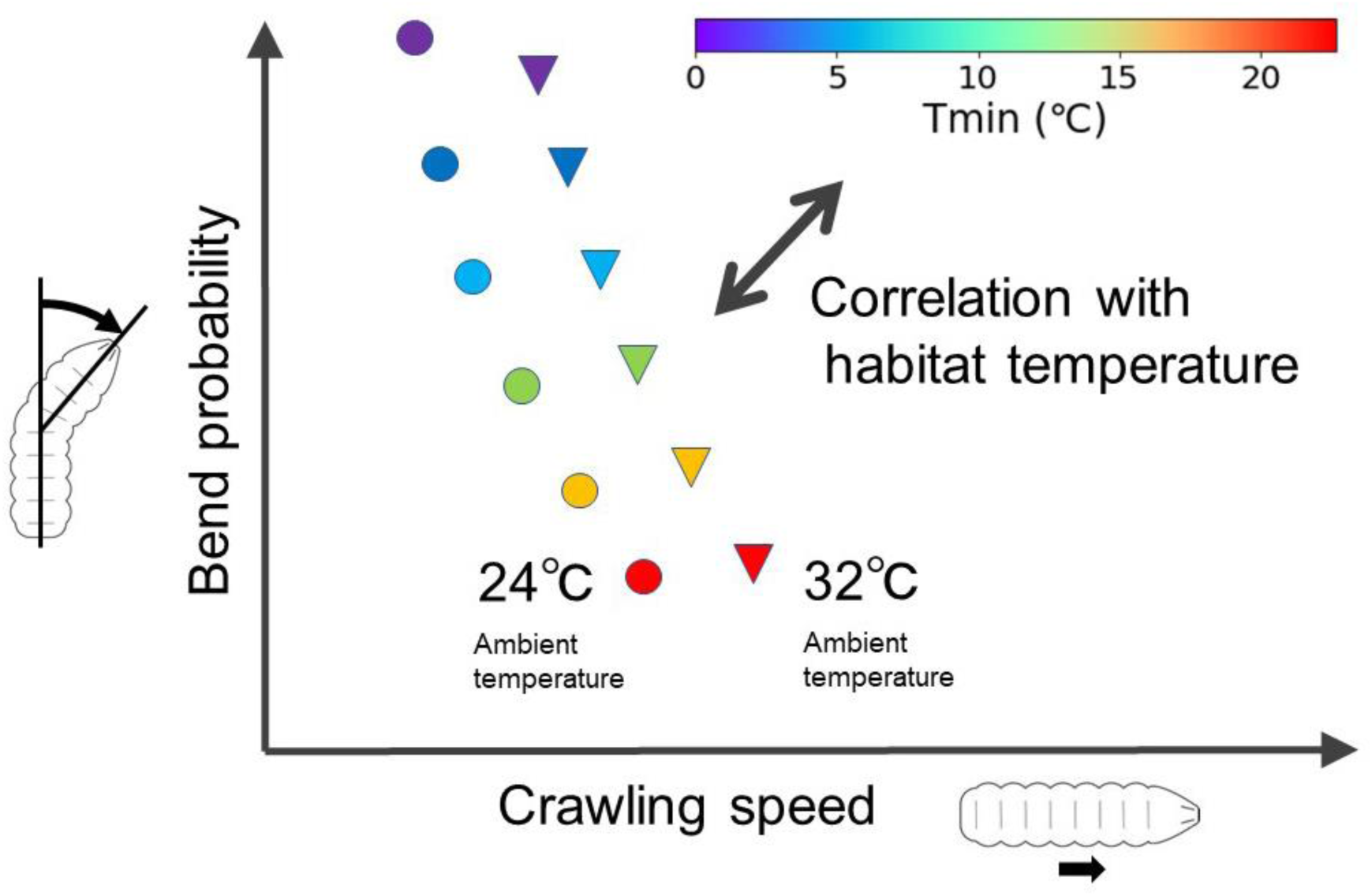
Summary of this study. Disks and triangles denote the kinematics of each species at 24°C and 32°C, respectively.

To conduct the quantitative analyses, we measured the larval crawling behaviour in simple experimental conditions: at the constant temperatures and humidity levels and on a flat surface of agarose gel. However, the environmental conditions in the wild for *Drosophila* larvae are far more complicated and diverse (Markow, 2015). For example, *Dmoj* breeds in cacti in the desert while *Dvir* inhabits in the slime fluxes in temperate and subarctic climate (Supplementary Figure 3; (Markow, 2015)). Our data in this paper imply that the variation of the kinematics should be related to the diversity of environmental conditions including habitat temperature. Comprehensive and quantitative analyses of larval locomotion in nature would shed light on the causes and meanings of the changes in larval kinematics appeared in the evolution.

The correlation between habitat temperature and the kinematics raises a question about the underlying mechanisms of adaptive change in kinematics to the habitat temperature. We presume that the adaptive change was likely to be driven by two factors: the pupation positioning and competition over food. Regarding the pupation behaviour, a previous study showed a correlation between path lengths of crawling and heights of pupation location in *D. melanogaster* (Sokolowski, 1985). Since pupae are immobile, adequate selection of pupation position is vital for their survival. Moisture is one of the key factors that affects the selection of pupation site, since dry environments can cause pupal desiccation, while soaked media might lead to the drowning of pupae (Sokolowski, 1985; Sokolowski et al., 1984). It is reasonable to assume that habitat temperature is a leading factor in determining the moisture levels of microenvironments of *Drosophila* larvae. Accordingly, divergent temperatures in the habitats of flies leads to variability in moisture levels, the ambient moisture affects the pupation position (Sokolowski et al., 1984), and the pupation position is related to the path length of crawling (Sokolowski, 1985). These three links might underlie the relationship between the larval locomotion kinematics and habitat temperature of each species.

The second possible mechanism underlying the relation between habitat temperature and locomotion kinematics is related to competition over food. It is suggested that larval feeding is competitive between individuals (Rohlfs & Hoffmeister, 2003). There is a positive relation between fitness and density (called Allee effects) in *D. melanogaster* larvae (Wertheim et al., 2002) and so female flies aggregate eggs when laying them (Rohlfs & Hoffmeister, 2003). Consequently, the larval population tends to be overcrowded (Burnet et al., 1977) and larvae compete over the limited resources to meet their need to consume sufficient amount of nutrient within limited larval periods (Nicholson, 1954). If the available food is limited and larval density is high, larvae need to crawl further (Sokolowski et al., 1997), which can be an evolutionary driving force to increase the crawling distance. In contrast, if there is plenty of food in the habitat, the crawling distance remains unchanged or even decreases during evolution, because crawling behaviour is energetically costly (Berrigan & Lighton, 1993). Consistent with this implication, the diversification in the larval kinematics in the 11 *Drosophila* species gives rise to the variation in the crawling distance (Figure 5). Accordingly, feeding conditions, especially the choice of which foods to eat, can affect kinematics of larval locomotion in evolutionary processes so that the divergence of food to eat can lead to the divergence in locomotion kinematics. In regard to this point, food for larvae of the genus *Drosophila* is divergent and related to their living environments (Markow, 2015; Watada et al., 2020; Watanabe et al., 2019). So, the nutritional conditions (nutrient balances and fermentation etc.) and physical properties (hardness and wetness etc.) of food to eat vary among the *Drosophila* species, which can drive evolutionary divergence in the kinematics of larval locomotion. Then, the growth of plants, including fruits and vegetables, for larvae is strongly affected by habitat temperature. Therefore, divergence in habitat temperature may affect the locomotion kinematics during evolution through divergence in the foods that larvae feed on, and divergence in requisite feeding behaviour for larval growth. Comprehensive kinematics studies of other *Drosophila* species and quantitative analyses of microenvironments of wild larvae in nature will give us insights on the relationships between ambient temperature and diverse larval locomotion kinematics.

How can we approach the neural circuit mechanisms underlying the interspecies divergence in larval locomotion? Circuit mechanisms in larval locomotion have been examined intensively in *Drosophila melanogaster*. Recent connectomics studies have identified several key neurons for larval locomotion in *Drosophila melanogaster* (Carreira-Rosario et al., 2018; M. Q. Clark et al., 2018; Fushiki et al., 2016; Heckscher et al., 2015; Hughes & Thomas, 2007; Kohsaka et al., 2014, 2017, 2019; Schneider-Mizell et al., 2016; Tastekin et al., 2015; Zwart et al., 2016). Regarding bending, the thoracic neuromere was shown to be important in bending in chemotaxis (Tastekin et al., 2015). Especially, the commissural connection is crucial for bending (Berni, 2015). In addition, a signal from the chordotonal sensory organ is also required to generate bending (Ainsley et al., 2003). Regarding the crawling speed, a group of inhibitory premotor neurons were identified to be critical (Kohsaka et al., 2014). Proprioceptive feedback and neuromodulation are both important for the normal crawling speed (Fox et al., 2006; Hughes & Thomas, 2007). Furthermore, kinematics of the larval locomotion have been measured and investigated quantitatively in detail (Gepner et al., 2015; Gershow et al., 2012; Gomez-Marin et al., 2011; Gomez-Marin & Louis, 2014; Heckscher et al., 2012; Hernandez-Nunez et al., 2015; Lahiri et al., 2011; Luo et al., 2010) and analysed by mathematical modelling (Gjorgjieva et al., 2013; Loveless et al., 2019; Loveless & Webb, 2018; Pehlevan et al., 2016; Sun et al., 2020). Regarding temperature sensing, neuronal and molecular mechanisms on temperature-guided behaviour have been clarified (Dillon et al., 2009; Klein et al., 2015; Kwon et al., 2008, 2010; Liu et al., 2003; Ni et al., 2016; Rosenzweig et al., 2005b, 2008; Shen et al., 2011). These extensive findings on the cellular and molecular mechanisms on larval crawling in *Drosophila melanogaster* will be an ideal starting point to investigate the evolution of larval behaviour in the genus *Drosophila*. For example, differences in commissure fibre tracts in the central nervous system (Berni, 2015) among the species might underlie the divergence in the bend probability. Interspecies comparison of a group of interneurons PMSIs (*period*-positive median segmental interneurons), which are involved in the crawling speed (Kohsaka et al., 2014), would reveal neural mechanisms on the evolutionary diversification in the crawling speed. Comparative analyses of neural network architectures and gene expression among the *Drosophila* species will relate the evolution of the nervous system to the adaptive diversification in larval behaviour.

## Materials and Methods

### Drosophila strains

We used the following fly stocks (17 strains from 11 species): *Drosophila ananassae* (k-s01), *Drosophila erecta* (k-s02), *Drosophila yakuba* (k-s03), *Drosophila melanogaster* (k-s04), *Drosophila sechellia* (k-s10), *Drosophila persimilis* (k-s11), *Drosophila pseudoobscura* (k-s12), *Drosophila willistoni* (k-s13), *Drosophila virilis* (k-s14) and *Drosophila mojavensis* (k-s15), *Drosophila melanogaster* collected at Kyoto (k-aba029) and Iriomote (k-aba032), *Drosophila ananassae* collected at Ogasawara (k-aaa027) and Iriomote (k-aaa309), and *Drosophila virilis* collected at Hokkaido (E-15601) and Toyama (E-15605) from KYORIN-Fly, Fly Stocks of Kyorin University and *Drosophila mauritiana* (#900020) from KYOTO Stock Center (DGRC) at the Kyoto Institute of Technology. These strains have been maintained at 23°C (except for *Drosophila persimilis* at 20°C) in the stock centres for more than 10 years. All animals were raised on standard cornmeal-based food at 25°C in the authors’ laboratory before the experiments for less than three months after the acquisition from the stock centres.

### Recording larval locomotion

Third-instar wandering larvae of each strain were picked up and gently washed in deionised water. Residual food on the larvae was brushed off with a paintbrush. An agarose stage (size: 9cm x 9cm x 5mm; 1.5% agarose; RIKAKEN STAR agarose powder #RSV-AGRP-100G) was placed on a temperature-controlled plate (Cool Plate, AS ONE Corporation, Japan) and the temperature of the surface was kept at 24±1°C, 32±1°C or 40±1°C. Eight to ten larvae were placed on the agarose arena with a paintbrush. The larvae were illuminated with infrared light, which larvae cannot see (LDQ-150IR2-850, wavelength = 850 nm, CCS Inc., Japan) and recorded at five frames/sec with a CCD camera (CGE-B013-U, MIGHTEX, Canada; resolution: 1280 pixels x 960 pixels) and an infrared filter (775LP filter, Omega Optical, USA), and saved as a series of bitmap files. In our setting, the scale of image is 0.13 mm/pixel. For each of the 17 strains (from 11 species) and at each of the three ambient temperatures, we repeated the measurement three times.

### Larva tracking

We obtained a time series of bitmap image files by the procedure described above. The bitmap files were converted to tiff format by Fiji (https://imagej.net/fiji). In cases where larvae accumulated at the first frame, we removed the first several images until the larvae dispersed enough to be identified individually. The x and y coordinates of the centroid and the bend angle (Figure 1E) of individual larvae were obtained by using the FIMTrack software (Risse et al., 2017). We used tracking data of single larvae that were continuously tracked for more than 300 frames (which corresponds to one minute). For the intraspecific analysis in Figure 4, single larvae with more than 150 continuous frames were tracked. In case a larva collided with another larva in the middle of the recording and the larva was traced differently before and after the collision with distinct labels, we treated the two traces as two larvae. Crawling speed of each larva was obtained as the median of crawling speed in the whole trace of the larva. Bend probability of each larva was calculated as the ratio of the number of frames showing bending (Figure 3 for the definition of bending) to the total number of the frames of its trace. The data of the three movies for each species/strain were merged. The coordinates and bend angles were smoothed and plotted by Python 3.7. For the visualization of the traces in Figure 1E and 1F, we smoothed the data by a uniform window function with five frames. The number of pixels in single larva ranges from about 150 (*Dyak*) to 500 (*Dpse*) that is sufficient to capture the body bend angles.

### Clustering analysis of kinematics

For clustering analysis of the plots in Figure 2A, we made a probability distribution. We discretised the centroid speed axis by an interval of 0.1 mm/sec between 0 and 4 mm/sec and the bend angle axis by an interval of two degrees between 0 and 140 degrees, then created 2800 bins in total (40 bins in the speed axis and 70 bins in the bend angle axis). For each scatter plot, we counted the number of points in each bin and obtained probability density by normalisation with the total number of the points. We calculated the Kullback-Leibler divergence (KL) of probability distributions of two species p(bin) and q(bin) as:

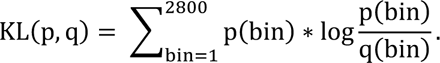

Since KL(p,q) is an asymmetric measure (KL(p, q) ≠ KL(q, p)), we calculated the Jensen-Shannon divergence JS, which is a symmetric alternative to the Kullback-Leibler divergence:

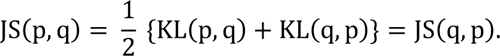

Based on the Jensen-Shannon divergence, we conducted the hierarchical clustering using Python 3.7 and the SciPy library.

### Habitat temperature of the *Drosophila* species

Global climate data were obtained from the WorldClim data website (https://www.worldclim.org/data/worldclim21.html). We used minimum, mean and maximum temperature in the world at a spatial resolution of five minutes (9.25 km x 9.25 km). Habitat regions of each *Drosophila* species were obtained from the literature (Brake & Baechli, 2008; Makino & Kawata, 2012; Markow & O’Grady, 2005) and the DrosWLD-Species database (https://bioinfo.museum.hokudai.ac.jp/db/modules/stdb/index.php?action=tbl&tbl_id=21) and traced them on the world map (2160 pixel x 4320 pixel) obtained from the WorldClim by using Adobe Illustrator software (Adobe Inc., USA). The maps in Supplementary Figure 2 were generated by Adobe Illustrator and the statistics of the temperature within each species’ habitat were calculated by Python 3.7.

### Statistical analysis

Statistical calculation was conducted by Python 3.7 in the following analyses: Kruskal-Walls test in Figure 3C and 3D, Pearson correlation in Figure 6B, 6C, 7B, 7C, 8B and 8C, Supplementary Figure 2B, 2C, 4B and 5B, and Mann-Whitney U-test in Figure 7A and 8A and Supplementary Figure 5A and 6A.

### Phylogenetic analyses

We adopted Bayesian statistics for phylogenetic analyses. In all the phylogenetic analyses, we used the software package, RevBayes v.1.0.7 (Hohna et al., 2016).

We estimated the *Drosophila* phylogenetic trees (Figure 8A) by using eight nuclear loci from Turelli et al. (2018): *aldolase*, *bicoid*, *enolase*, *esc*, *transaldolase*, *white*, *wingless*, and *yellow*. We used *Scaptodrosophila lebanonensis* (*S. lebanonensis*) as an outgroup. Coding sequences of these genes for all the species except for *D. mauritiana* and *S. lebanonensis* were obtained from Kalay et al., (2020). Coding sequences of the genes of *D. mauritiana* and *S. lebanonensis* were obtained from the NCBI web site (https://www.ncbi.nlm.nih.gov/). The genes were aligned with MAFFT version 7 (Katoh & Standley, 2013). We selected the eight genes out of twenty genes described in Turelli et al., (2018) because homologous genes of the other twelve genes were not identified in the genome assembly of *D. mauritiana*. Based on the eight coding sequences of the twelve species, we performed chronogram analyses described in Turelli et al. (2018) and obtained the phylogenetic tree in Figure 8A, where the root is the branch of *S. lebanonensis* and the length of each branch indicates relative time in the evolution. We repeated the phylogeny inference four times to obtain four trees to check the robustness of this inference in the following analysis.

Based on the phylogenetic tree obtained above, we analyzed the evolution of kinematics parameters. We used eight parameters (bend probability at 24, 32 and 40°C, crawling speed at 24, 32 and 40°C, backward crawling probability at 40°C, and body length), for each of the 11 *Drosophila* species. In this analysis, while the topology of the phylogenetic tree is unchanged, the rates of evolution of the kinematic parameters and the rates of evolution of the branches of the phylogenetic tree were estimated by Bayesian inference. We assumed the kinematics parameters evolve under a multivariate Brownian-motion model (Huelsenbeck & Bruce, 2003; Lartillot & Poujol, 2011). This model consists of two components: the relative rates of evolution among the kinematics parameters and the correlation between each pair of the kinematics parameters (Kalay et al., 2020). The Brownian model assumes that the changes in the kinematics parameters during evolution are additive, which has two consequences: 1) parameters can become negative; 2) evolution rates can depend on the scale of the kinematics parameters. To cope with these properties, we adopted log-transformation to the kinematics parameters as described previously (Kalay et al., 2020). This transformation guarantees that the original kinematics parameters (which are exponentials of the log-transformation) remain positive and controls the size issue by transforming additive changes to multiplicative ones (Kalay et al., 2020). For prior models for the evolution rates of the branches, we specified an uncorrelated exponential (UCED) relaxed-clock model, where the rate of each branch is drawn independently from an exponential distribution. We ran the Markov Chain Monte Carlo (MCMC) simulation of UCED with each of the four phylogenetic trees described in the previous paragraph (Figure 11). Consistent results were obtained by another prior model, an uncorrelated gamma (UCG) relaxed clock model (Supplementary Figure 7), which indicates the predictions are robust to the choice of priors.

## Acknowledgement

We thank KYORIN-Fly, Fly Stocks of Kyorin University and KYOTO Stock Center (DGRC) at the Kyoto Institute of Technology for providing fly lines, Dr. Takashi Makino for advice about climate data in the world. We thank Ms. Shibahara for tracing habitats of the 11 species. We thank Dr. Masayoshi Watada for critical comments on the paper. This work was supported by MEXT/JSPS KAKENHI grants (17K19439, 19H04742, 20H05048 to A.N. and 17K07042, 20K06908 to H.K.).

## Competing interests

We have no conflicts of interest with respect to the work.

## Notes

### Competing Interest Statement

The authors have declared no competing interest.

