## Supplemental Figures for "Interspecies variation of larval locomotion kinematics in the genus *Drosophila* and its relation to habitat temperature"

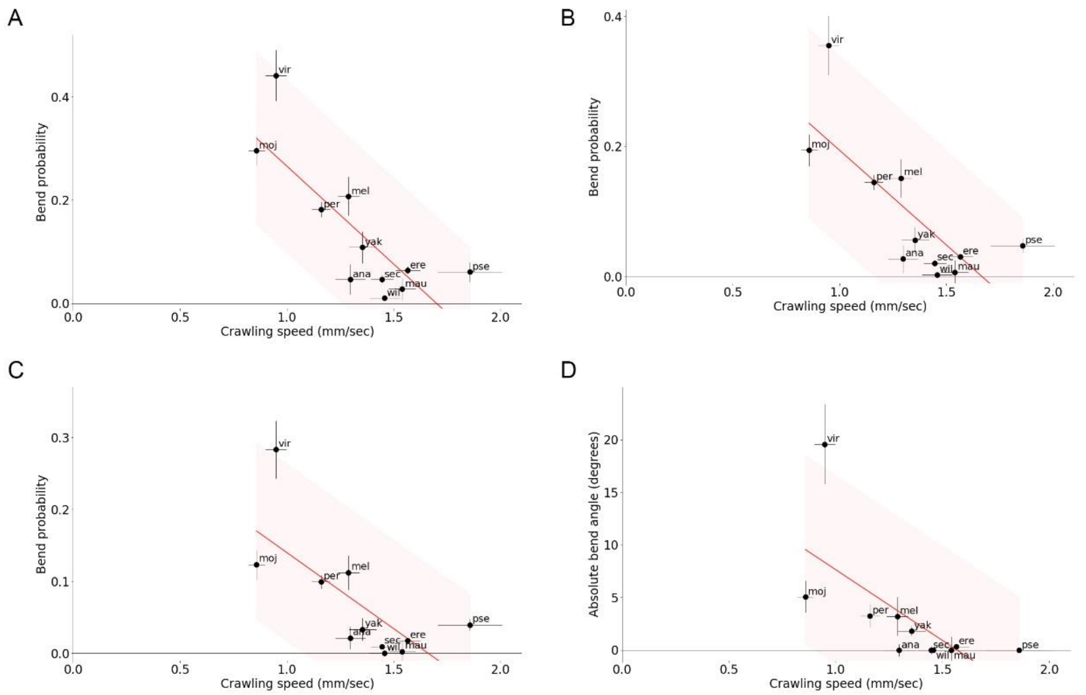


**Supplementary Figure 1. The negative correlation between the crawling speed and bend probability is robust to the angle threshold for the definition of bend.** (A-C) Scatter plots of the bend probability at 24°C against the crawling speed at 24°C. (A) The angle threshold for bend is 10 degrees. (B) The threshold is 20 degrees. (C) The threshold is 30 degrees. Median ± sem is shown. B is the same panel as Figure 3C represented here for comparison. (D) Scatter plot of the median of absolute bend angle at 24°C against the crawling speed at 24°C. The red lines show the linear regression functions and the shaded areas represent the 95% confidence band. The point estimates of the Pearson correlation and its 95% confidence intervals are -0.80 and [-0.95, -0.39] in A, -0.76 and [-0.94, -0.30] in B, -0.71 and [-0.92, -0.19] in C, and -0.66 and [-0.90, -0.11] in D.

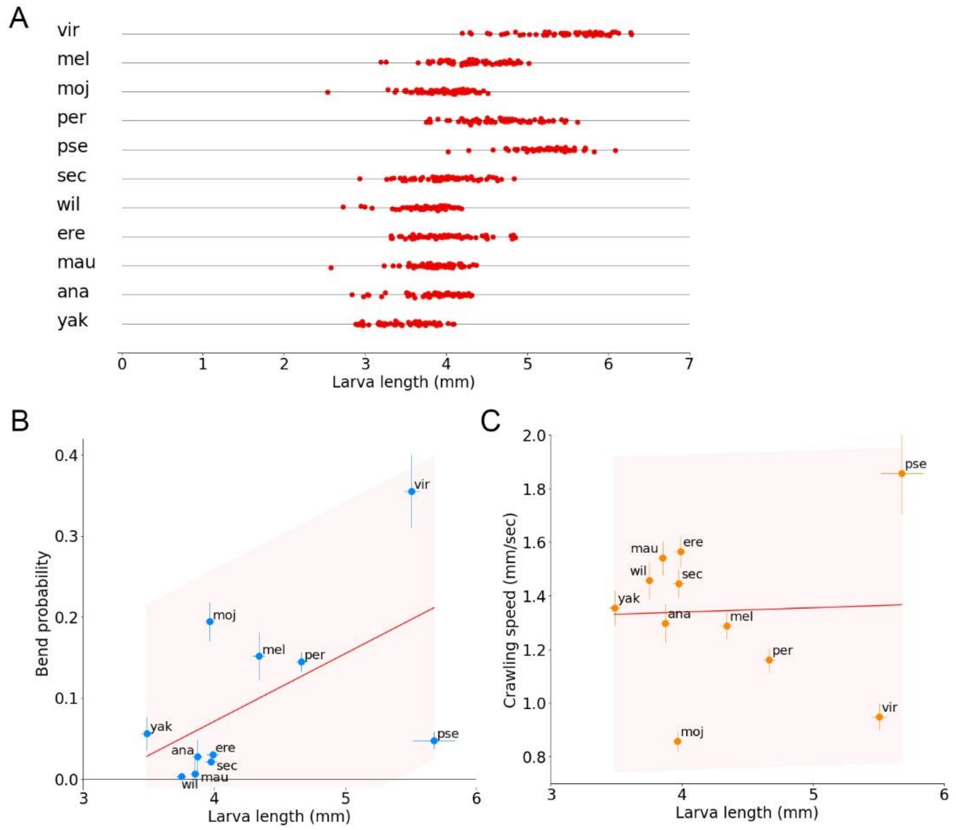


**Supplementary Figure 2. Interspecies comparison between larva length and kinematics parameters.** (A) Axial body length of larvae of the 11 species. Sample numbers are the following: *Dvir*: n=69; *Dmel*: n=72; *Dmoj*: n=82; *Dper*: n=76; *Dpse*: n=60; *Dsec*: n=68; *Dwil*: n=66; *Dere*: n=69; *Dmau*: n=69; *Dana*: n=68; *Dyak*: n=56. (B) Scatter plot of the bend probability at 24°C against the larva length of the 11 species. (C) Scatter plot of the crawling speed at 24°C against the larva length of the 11 species. In B and C, median ± sem are shown. The red lines show the linear regression functions and the shaded areas represent the 95% confidence band. The point estimates of the Pearson correlation and its 95% confidence intervals are 0.55 and [-0.07, 0.87] in B and 0.04 and [-0.57, 0.63] in C.

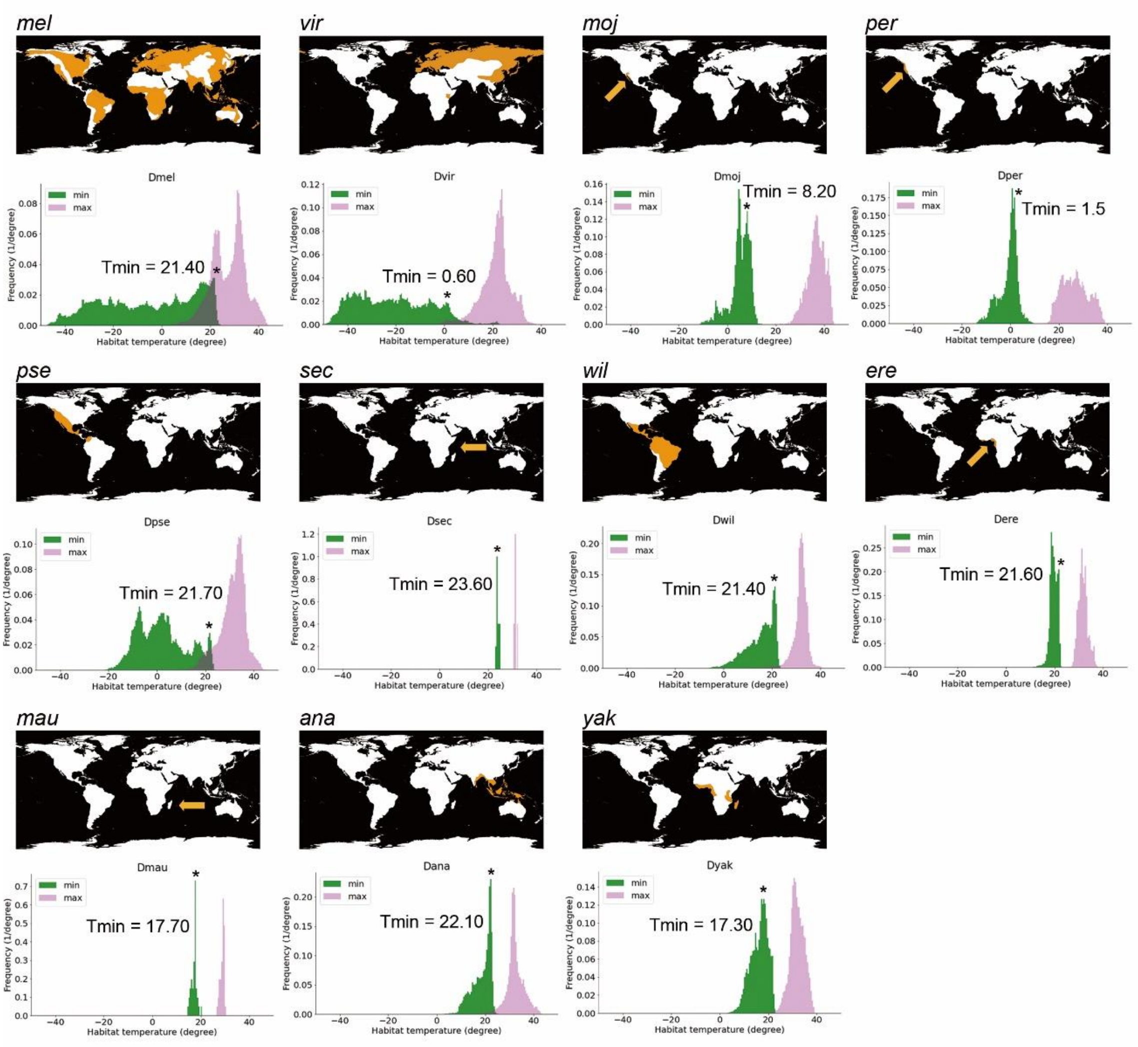


**Supplementary Figure 3. World maps of histograms of habitat temperature of the *Drosophila* species.** The top of each panel shows a map of the habitat of the labelled species. The bottom on each panel shows a histogram of minimum (green) and maximum (magenta) temperatures within the habitat. Asterisks denote the warmest peak in the minimum habitat temperature. The temperature at the peak is called Tmin in this study.

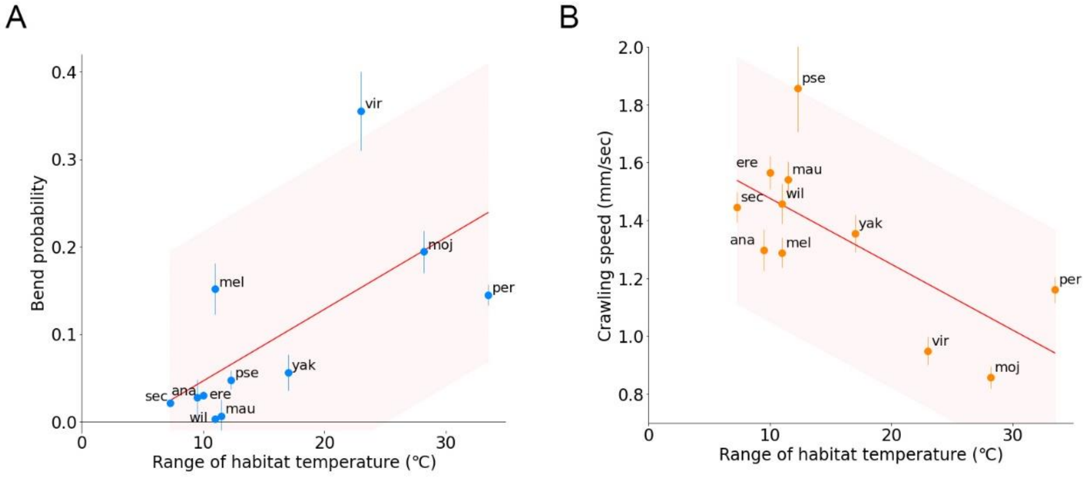


**Supplementary Figure 4. Relationship between the kinematics of larval locomotion and the range of habitat temperature of the Drosophila species** (A) Scatter plot of the bend probability speed at 24°C against the range of habitat temperature of the 11 species. (B) Scatter plot of the crawling speed probability at 24°C against the larva length of the 11 species. Median ± sem are shown. The red lines show the linear regression functions and the shaded areas represent the 95% confidence band. The point estimates of the Pearson correlation and its 95% confidence intervals are 0.65 and [0.08, 0.90] in A and -0.69 and [-0.91, -0.15] in B.

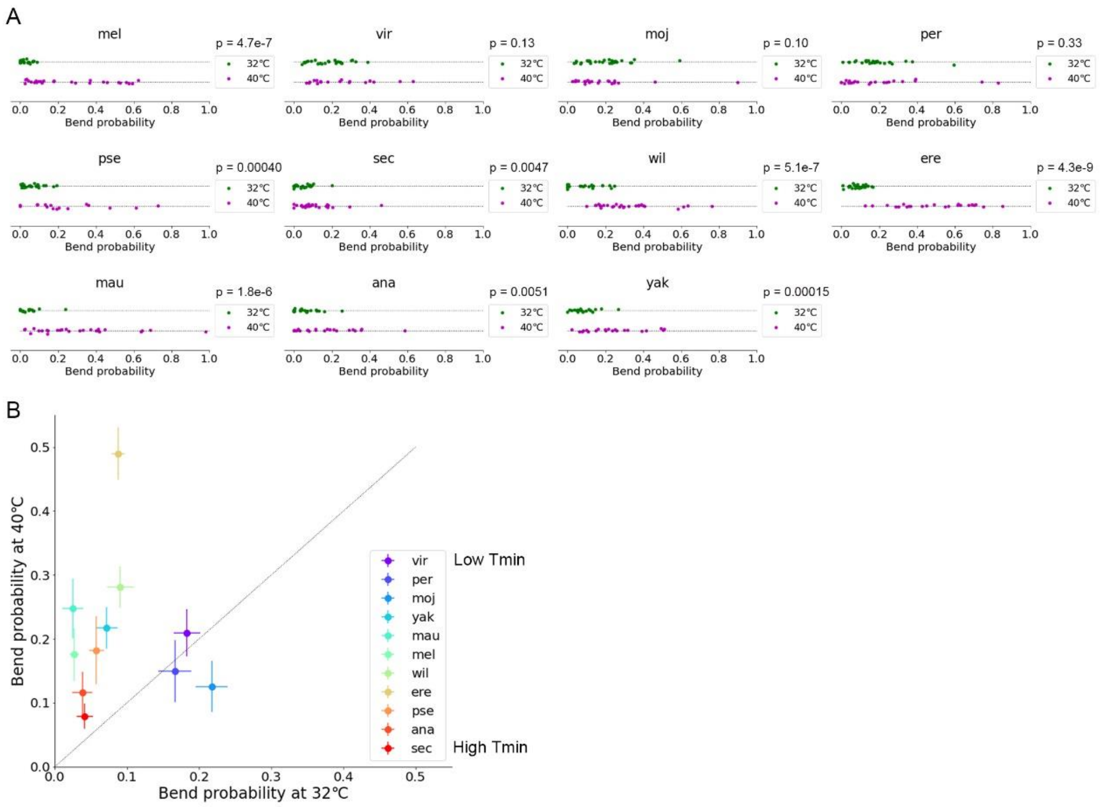


**Supplementary Figure 5. Bend probability in the 11 *Drosophila* species at 40°C.** (A) Bend probability of the 11 species at 32°C and 40°C. The data at 32°C are the same as in Figure 4A. (B) Scatter plot of the bend probability at 40°C against at 32°C. Sample numbers at 40°C are the following: *Dvir*: n=20; *Dmel*: n=28; *Dmoj*: n=23; *Dper*: n=21; *Dpse*: n=16; *Dsec*: n=27; *Dwil*: n=27; *Dere*: n=22; *Dmau*: n=26; *Dana*: n=23; *Dyak*: n=20. Sample numbers at 32°C are described in the legend of Figure 4. Median ± sem are shown in B.

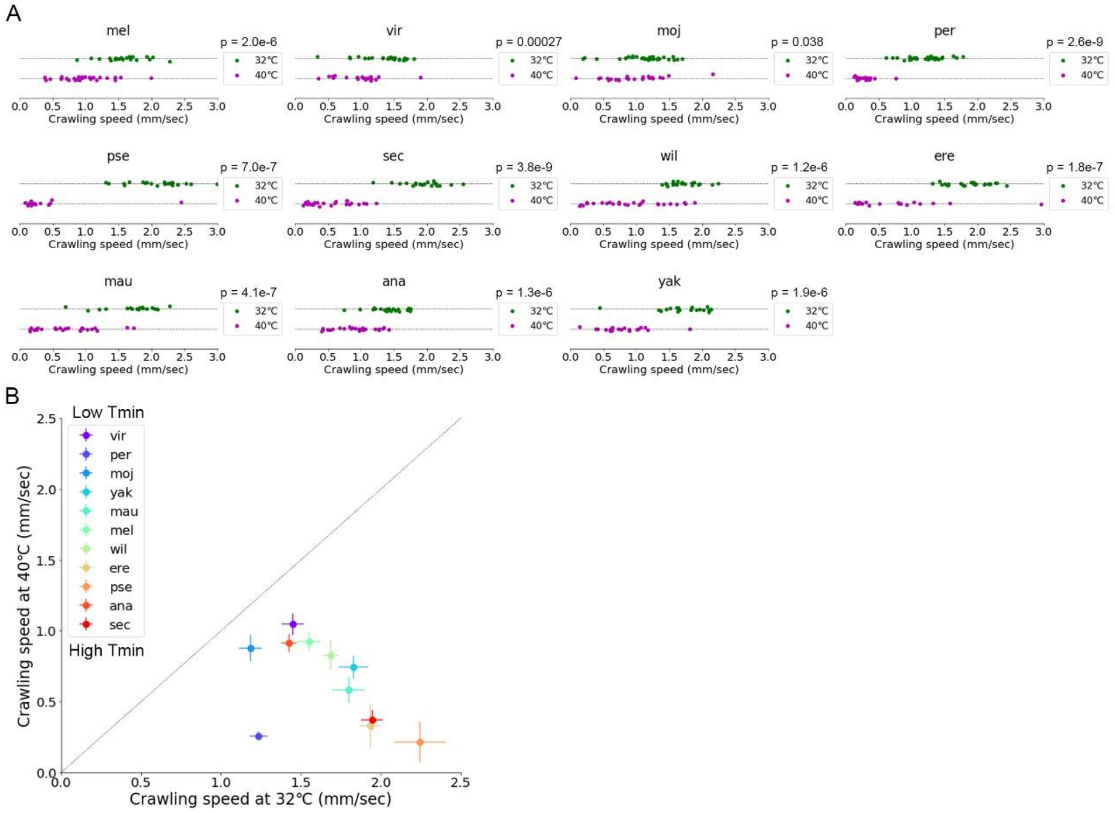


**Supplementary Figure 6. Crawling speed in the 11 *Drosophila* species at 40°C.** (A) Crawling speed of the 11 species at 32°C and 40°C. The data at 32°C are the same as in Figure 5A. (B) Scatter plot of the crawling speed at 40°C against at 32°C. Sample numbers at 40°C are the following: *Dvir*: n=20; *Dmel*: n=28; *Dmoj*: n=23; *Dper*: n=21; *Dpse*: n=16; *Dsec*: n=27; *Dwil*: n=27; *Dere*: n=22; *Dmau*: n=26; *Dana*: n=23; *Dyak*: n=20. Sample numbers at 32°C are described in the legend of Figure 5. Median ± sem are shown in B.

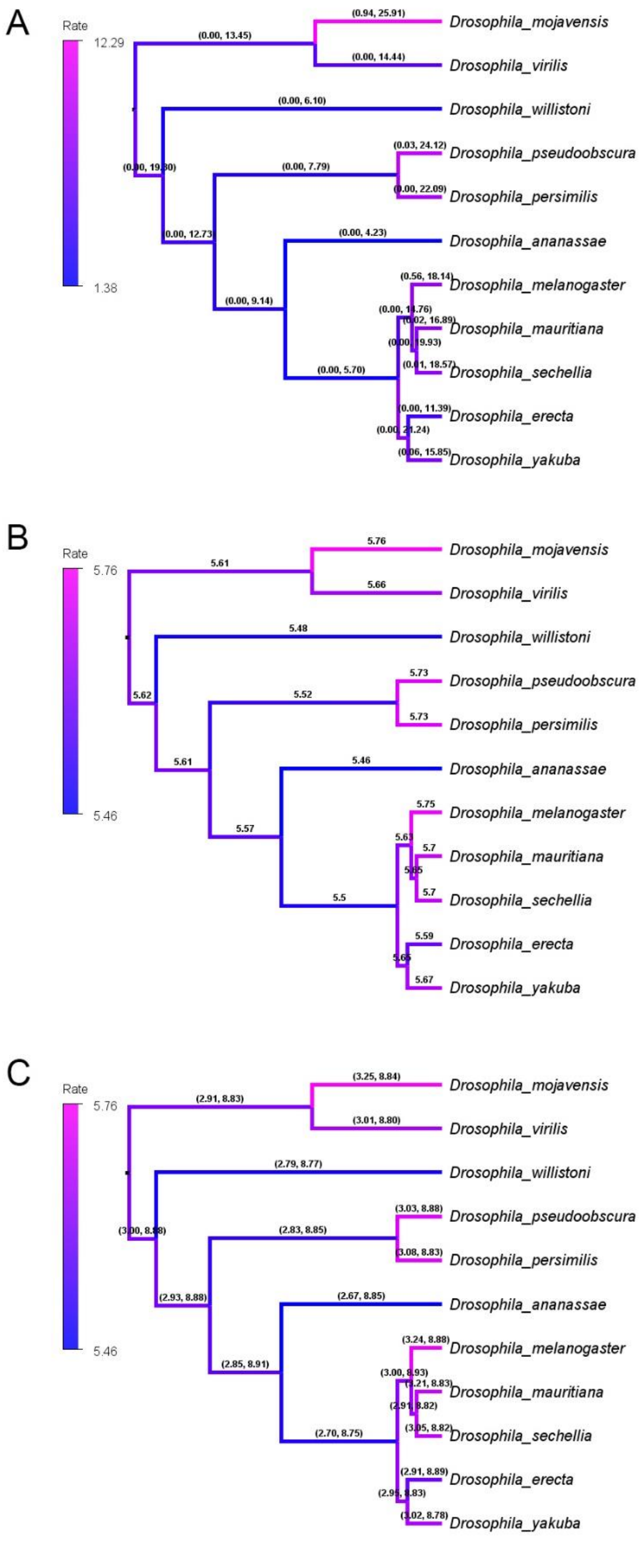


**Supplementary Figure 7. Phylogenetic analyses of larval locomotion based on distinct**

**prior models.** (A) Phylogenetic tree estimated from Bayesian inference with an uncorrelated exponential (UCED) relaxed-clock model as the prior. Values of each branch denote the range of 95% highest posterior density of the inference of the evolution rates shown in Figure 11A. (B and C) Phylogenetic tree estimated from Bayesian inference with an uncorrelated gamma (UCG) relaxed clock model. Values of each branch show the evolution rate (B) and its range of 95% highest posterior density (C).
